# Comparison of dental topography of marmosets and tamarins (Callitrichidae) to other platyrrhine primates using a novel freeware pipeline

**DOI:** 10.1101/2023.08.31.555703

**Authors:** Dorien de Vries, Mareike C. Janiak, Romina Batista, Jean P. Boubli, Ian B. Goodhead, Emma Ridgway, Doug M. Boyer, Elizabeth St. Clair, Robin M.D. Beck

## Abstract

Dental topographic metrics (DTMs), which quantify different aspects of the shape of teeth, are powerful tools for studying dietary adaptation and evolution in mammals. However, comparative samples of scanned mammal teeth suitable for analysis with DTMs remain limited in size and scope, with little or no representation of some major lineages, even within well-studied clades such as primates. In addition, current DTM protocols usually rely on proprietary software, which may be unavailable to many researchers for reasons of cost. We address these issues in the context of a DTM analysis of the primate clade Platyrrhini (“New World monkeys”) by: 1) presenting a large comparative sample of scanned second lower molars (m2s) of callitrichids (marmosets and tamarins), which were previously underrepresented in publicly available platyrrhine datasets; and 2) giving full details of an entirely freeware pipeline for DTM analysis. We also present an updated discrete dietary classification scheme for extant platyrrhines, based on cluster analysis of dietary data extracted from 98 primary studies. Our freeware pipeline performs equally well in dietary classification accuracy of an existing sample of platyrrhine m2s (excluding callitrichids) as a published protocol that uses proprietary software, at least when multiple DTMs are combined. Individual DTMs however, sometimes showed very different results in classification accuracies between that of our freeware pipeline and that of the proprietary protocol, most likely due to the differences in the smoothing functions used. The addition of callitrichids still resulted in high classification accuracy in predicting diet with combined DTMs, although accuracy was considerably higher when molar size was included (90%) than excluded (73%). We conclude that our new freeware DTM pipeline is capable of accurately predicting diet in platyrrhines based on tooth shape and size, and so is suitable for inferring probable diet of taxa for which direct dietary information is unavailable, such as fossil species.

## Introduction

Dental topographical metrics (DTMs) attempt to quantify functional and adaptive aspects of tooth shape, with different dental topographic metrics capturing different functional aspects (Kay 1975; Kay and Hylander 1978; Strait 1993a; b; Zuccotti et al. 1998; Ungar and M’Kirera 2003; Cuozzo and Sauther 2006; Evans et al. 2007; Evans and Jernvall 2009; Bunn et al. 2011; Guy et al. 2013; Tiphaine et al. 2013; Berthaume et al. 2018, 2019a). DTMs are increasingly used for studying the relationship between dental morphology and diet. Shearing Quotient (Kay 1975), and Shearing Ratio (Strait,1993a; b) were some of the first DTMs to be proposed. These metrics capture two-dimensional shape of teeth and require identification of homologous landmarks on the occlusal surface of the tooth. Newer DTMs capture three-dimensional tooth shape and can be computed with minimal reference to specific morphological features (and thus are often said to be “homology free”, Evans et al. 2007), making them useful for meaningfully comparing tooth shape between taxa that may lack clearly homologous structures (Evans et al. 2007; Evans and Jernvall 2009; Prufrock et al. 2016).

DTMs have been particularly widely used to compare tooth shape in primates. These comparisons have been successfully employed to characterise and distinguish different dietary groups of many clades of extant primates, based on analyses of upper molars (e.g., Ungar et al. 2018), lower molars (e.g., Boyer 2008; Bunn and Ungar 2009; Bunn et al. 2011; Winchester et al. 2014; Berthaume and Schroer 2017), or both together (e.g., Allen et al. 2015). DTMs have also been used to predict the diets of extinct primates (see Table 1 of Berthaume and Schroer 2017 for an overview of studies). Reliable dietary predictions require an appropriate comparative sample of extant taxa that exhibit a diverse range of known diets that includes those likely to have been present in the extinct species (Berthaume and Schroer 2017). Ideally, the comparative extant dataset should also take into account the phylogenetic relationships of the fossil taxa to be tested, as the same dietary category can be reflected by different dental topographic values in different clades, as shown by Winchester et al. (2014) using a sample of platyrrhines, strepsirrhines, and tarsiers. Thus, although new DTMs are homology-free, they are not “phylogeny free”.

**Table 1.**
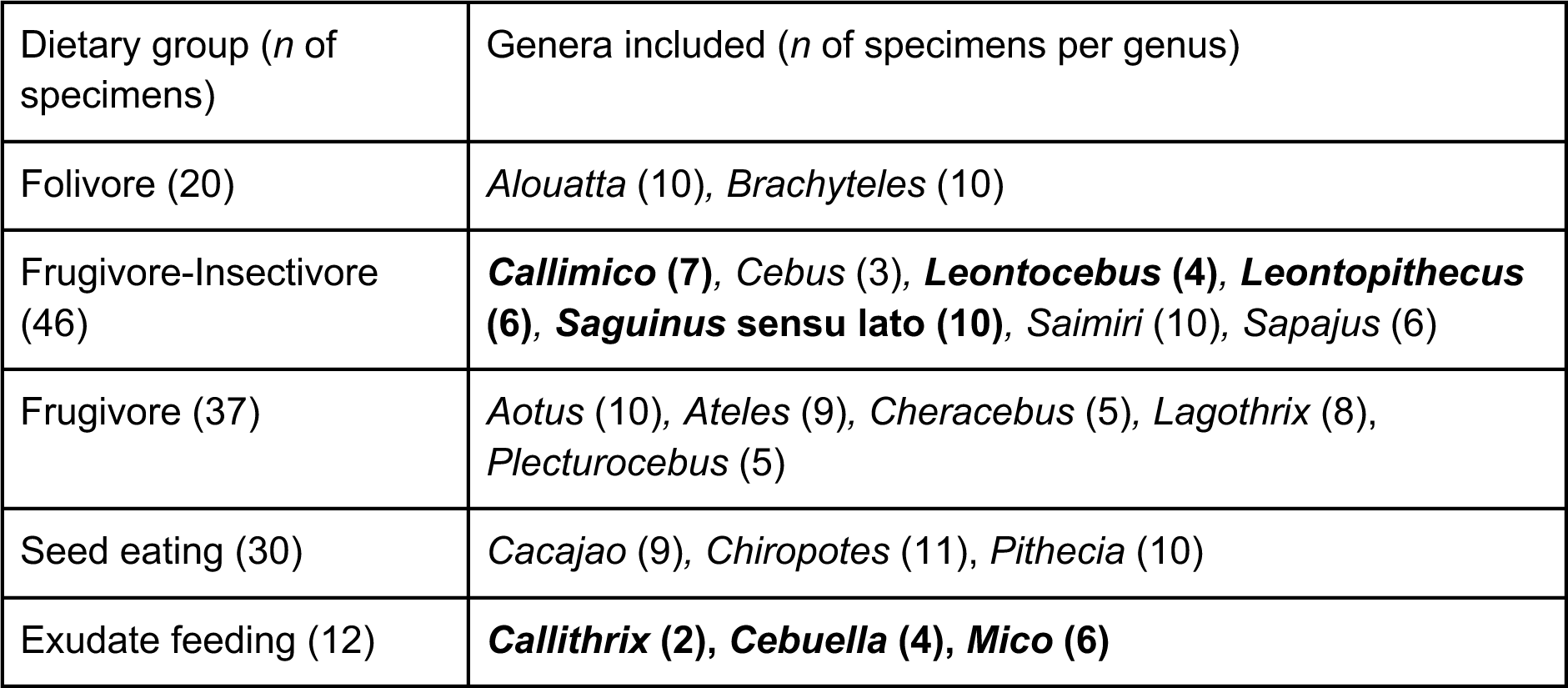
Sample size breakdown of the entire sample (n=145) per dietary category following the preferred dietary scheme resulting from this study (edited-UPGMA scheme). The newly added callitrichid specimens (n=39) are listed in bold

Although dental topographic methods represent a powerful, quantitative approach for analysing tooth shape, a number of issues currently limit their applicability. Firstly, current pipelines for dental topographic methods typically use commercial software packages that are often relatively expensive, and hence unaffordable for many researchers (but see Morley and Berthaume 2023). Secondly, studies that attempt to link tooth shape to particular diets often use dietary classification schemes that are not based on the full range of primary data available in the scientific literature. Thirdly, surface meshes suitable for dental topographic analysis (or scan data that can be used to generate these) are still only available for small subsets of known mammalian diversity, and hence studies using such data are typically quite limited in taxonomic scope. Even within primates there are gaps in data. They are especially limited with respect to the smallest platyrrhines, callitrichids. Here we address all three of these issues in relation to the primate clade Platyrrhini, as follows.

### A freeware pipeline for dental topographic analyses

Recent developments in dental topographic freeware have made methods for calculating dental topographic variables increasingly easily accessible and easy to use (R-package molaR, Pampush et al. 2016, 2022; freeware MorphoTester, Winchester 2016). However, until now most dental topographic protocols (e.g., Boyer 2008; Spradley et al. 2017; Fulwood et al. 2021; Pampush et al. 2022) have used proprietary software, such as Amira/Avizo and GeoMagic, for processing of raw scan data and digital surface meshes into the correct format for calculating dental topography (i.e., all specimens are consistently simplified to the same number of polygons, oriented into occlusal view along the z-axis, smoothed, and exported as a .ply file). Although some studies have mentioned that some steps can also be performed in the freeware package MeshLab (Winchester 2016; Melstrom 2017; Spradley et al. 2017; Berthaume et al. 2020), to our knowledge, an only one processing pipeline that uses free software throughout has been published and validated before (Morley and Berthaume 2023). The goal of Morley and Berthaume (2023) was to replicate the decimation and smoothing steps done in Avizo as closely as possible using freeware, whereas the aim of our freeware pipeline is to yield meaningful surface meshes suitable for distinguishing different diets using DTMs. To maximise the utility of this study to other researchers, we therefore present and validate a novel free pipeline for processing scan data for dental topographic analyses that uses only freeware, specifically: Slicer (Kikinis et al. 2014), the SlicerMorph extension (Rolfe et al. 2021), MeshLab (Cignoni et al. 2008), Autodesk MeshMixer (MeshMixer RRID:SCR_015736), R (R Core Team 2022), and the R-package molaR (Pampush et al. 2016).

### An improved dietary classification scheme for platyrrhine primates

There is a long history of studies that have applied formal dietary classification schemes for mammals (e.g., Harrison 1962; Andrews et al. 1979; Eisenberg 1981). In general, such schemes have used discrete categories, e.g., carnivore, insectivore, omnivore (but see Wisniewski et al. 2022, for an ordinal ranking-based approach). A number of different dietary schemes have been used in previous dental topographic studies of primates, and these are often based on primary data (e.g., stomach contents, behavioural observations), basing categories on the primary food source and a relatively limited range of studies (Boyer 2008; Cooke 2011; and sometimes secondary food source as well: Allen et al. 2015). In contrast to this, some recent studies have used clearly defined, quantitative approaches for dietary classification, based on detailed analysis of extensive primary scientific literature, notably Pineda-Munoz and Alroy (2014) and Lintulaakso et al. (2023). Here, we use cluster analysis of quantitative dietary data from 98 primary studies (the largest collection for a study of this kind) to produce a revised set of discrete diet categories for the extant primate clade Platyrrhini. Our dataset includes 20 of the 23 genera (= 87%) currently recognised within platyrrhines (IUCN SSC Primate Specialist Group).

### New dental topographic data for callitrichids

The clade Platyrrhini (variously referred to as “New World primates”, “monkeys of the Americas”, or “Neotropical primates”) has high extant taxonomic diversity, with 187 species in 23 genera. The crown platyrrhine radiation comprises as many as five families, depending on the classification used: Atelidae (howler monkeys, spider monkeys, and relatives), Pitheciidae (titis, sakis, and uakaris), Cebidae (squirrel monkeys and capuchins), Callitrichidae (marmosets and tamarins), and Aotidae (night or owl monkeys) (some place callitrichids as a subfamily within Cebidae, Harris et al. 2014; Kay 2015; Rosenberger 2020). Collectively, extant platyrrhines span a broad range of ecological niches, including a wide variety of diets and body sizes (but see Fleagle and Reed 1996; e.g., Norconk et al. 2009; Youlatos 2018). Callitrichids are the smallest extant members of Anthropoidea (= monkeys and apes), with the smallest species, *Cebuella pygmaea*, having an average body mass of 110-122 g (Smith and Jungers 1997). Extant callitrichids comprise tamarins (*Saguinus* spp.*, Tamarinus* spp.*, Oedipomidas* spp., *Leontocebus* spp.), lion tamarins (*Leontopithecus* spp.), marmosets (*Callithrix* spp., *Cebuella* spp., *Mico* spp., *Callibella humilis*), and Goeldi’s monkey (*Callimico*, IUCN SSC Primate Specialist Group). Callitrichids exhibit a range of diverse diets, and are known to consume insects, fruits, and fungi (Rylands and Faria 1993; Meireles et al. 1999; Harris et al. 2014). Marmosets (*Callithrix, Cebuella, Mico*, and *Callibella*) also display adaptations for feeding on exudates, which is unique as a primary dietary specialisation amongst anthropoids (although some strepsirrhines, such as *Euoticus* and *Phaner*, also regularly feed on exudates; Nash 1986).

Callitrichids are of particular importance for understanding platyrrhine dental and dietary evolution, and for reconstructing the diets of extinct platyrrhines, because the earliest known fossil platyrrhines, as well as some later taxa, were of similar size based on dental dimensions (Bond et al. 2015 p. 538; Antoine et al. 2016, 2017; Marivaux et al. 2016; Kay et al. 2019 p. 1). In particular, the oldest known probable platyrrhine, *Perupithecus ucayaliensis*, has been specifically likened to callitrichids in some features of its dental morphology (Bond et al. 2015). However, comparative datasets publicly available to study platyrrhine dental shape, such as that of 111 second lower molars created by Winchester et al. (2014 publicly available as MorphoSource Project ID 000000C89), suffer from a general lack of callitrichids. In fact, to our knowledge, there has been only one published study using dental topography that includes a limited number of species sampled for callitrichids (*Callithrix* and *Saguinus* sensu lato) amongst a broad platyrrhine sample of upper second molars (Ungar et al. 2018). A study on dietary adaptations and dental morphology by Kay et al. (2019) included a wide range of callitrichids, but used only one DTM (2D shearing quotient) as well as methods other than dental topography (3D GM landmark analysis). In addition, both Ungar et al. (2018) and Kay et al. (2019) used upper molars only; there are currently no dental topographic comparative studies of platyrrhine lower molars that include callitrichids, even though lower molars are more commonly used in such studies have been shown to be consistently more successful at predicting diet than uppers within non-callitrichid platyrrhines (Allen et al. 2015). The inclusion of callitrichids expands the range of diets and body sizes present in comparative datasets of tooth shape in Platyrrhini, increasing their usefulness for studying dietary evolution in this ecologically diverse and evolutionarily successful primate clade.

### Aims of study

Based on the above considerations, the aims of this paper are fourfold: 1) introduce and validate processing pipeline for dental topographic analyses solely using freeware; 2) present a scheme of dietary categories for extant platyrrhines based on all quantified components of their diet from a large dataset of dietary observations taken directly from the primary literature; 3) expand the publicly available sample of surface meshes of platyrrhine molars via the addition of 39 callitrichid m2 specimens, representing 7 extant callitrichid genera; 4), use our freeware pipeline and newly generated data to test the classification accuracy using the newly designed dietary scheme on the total (i.e. callitrichid and non-callitrichid) platyrrhine sample of surface meshes. The latter will be particularly useful for future studies reconstructing the diets of fossil platyrrhine taxa.

## Materials & Methods

### Platyrrhine sample

See Table 1 for the breakdown of the total sample per dietary category and per genus, see Supplementary Information Table S2 for specimen information. For our non-callitrichid platyrrhine sample of surface meshes, we downloaded 111 cropped but unsmoothed surface meshes from Winchester et al. (2014), which are available as project ID 000000C89 on MorphoSource.org (Boyer et al. 2016).

For the new callitrichid sample (see Figure 1), high-resolution plastic replica casts were made of 39 callitrichid lower second mandibular molars representing 10 species and 7 genera (see Supplementary Information Table S2 for full details) following the protocols of Boyer (2008) and Winchester et al. (2014). The callitrichid casts were scanned with a Scanco Medical brand μCT 40 scanner at 8 μm (for *Cebuella pygmaea* specimens) and 10 μm (all other callitrichid specimens). Surface meshes of the callitrichid sample were generated and cropped in Avizo. However, our expanded freeware protocol in the Supplementary Information includes instructions for performing this step using the freeware Slicer (Kikinis et al. 2014) and the SlicerMorph expansion (Rolfe et al. 2021). All further processing steps followed those of the freeware pipeline outlined below.

**Fig. 1.**
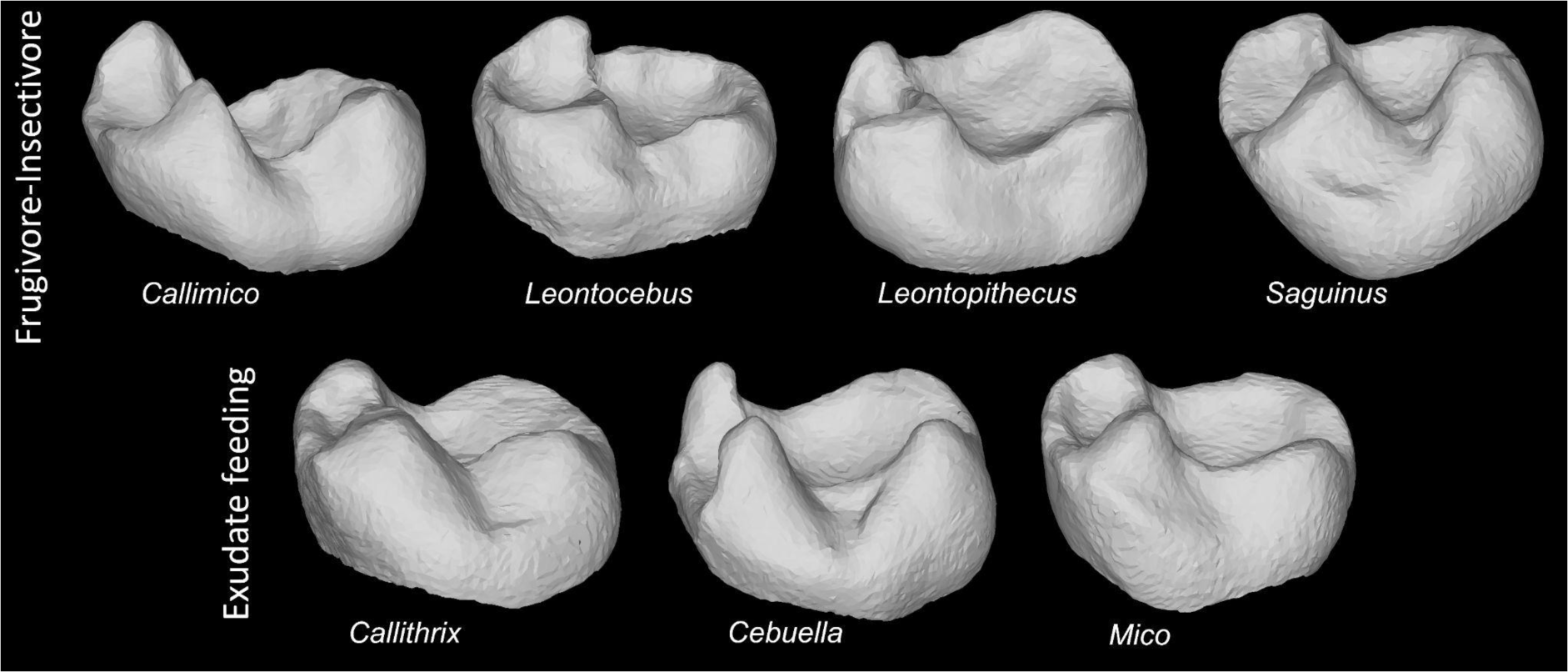
Examples of callitrichid lower second molar (m2) surface files processed using the described freeware pipeline for every genus of our sample. Teeth in the upper row belong to genera that are classified as frugivore-insectivore in our preferred dietary classification scheme (“edited-UPGMA), whilst those in the lower row belong to genera classified as exudate feeding in the same scheme. Images are not to scale

### Freeware pipeline for dental topographic analyses

To validate our freeware pipeline, we replicated the protocol of Winchester et al. (following Winchester et al. 2014) as closely as possible using freeware instead of commercial or proprietary software on the exact same non-callitrichid platyrrhine sample only. After this validation step, the new callitrichid specimens were processed using the same freeware protocol.

### Surface mesh processing

The outlined surface mesh processing steps can all be performed on an up-to-date standard laptop (4×4GB RAM), which provides sufficient computational capabilities. The surface meshes from the Winchester et al. (2014) sample needed to be flipped along the z-axis to be oriented correctly, which was done in MeshLab (Cignoni et al. 2008) using a custom MeshLab script written by the first author (see Supplementary Information).

First, all surface meshes were manually inspected in Meshmixer v3.5.474 (MeshMixer RRID:SCR_015736), with any deformities such as small cracks in the enamel or bubbles introduced during the moulding process manually reconstructed, if necessary. Then, using MeshLab v2022.2 (Cignoni et al. 2008), the reconstructed surface meshes were centred, oriented into occlusal view, cleaned of abnormalities (e.g., removing duplicate faces, removing isolated pieces), downsampled to 10,000 triangles (following Winchester et al. 2014; Spradley et al. 2017; Berthaume et al. 2019b), and smoothed using four different smoothing settings.

Unlike Amira/Avizo, MeshLab offers various different smoothing options, all altering the surface mesh in different ways. We explored four different smoothing options: the ‘HC Laplacian Smooth’ (HCL) option (Vollmer et al. 1999), and the ‘Taubin Smooth’ option set to 10 (TAU10), 50 (TAU50), and 100 (TAU100) iterations (Taubin 1995). Based on qualitative visual inspection, the outputs of these smoothing settings were visually closest to the results of smoothing a surface mesh in Amira/Avizo, and, importantly, they did not reduce the 3D surface area of the surface mesh (also identified by Hafez and Rashid 2023) unlike other smoothing functions in MeshLab. The chosen four smoothing options differed in the degree of the strength of smoothing, with HCL being the lightest smoothing setting, and TAU100 being the strongest smoothing setting. As Spradley et al. (2017) showed that excessive smoothing can create sharp, horn-like artefacts on the surface mesh, surface meshes were inspected after smoothing to make sure no artefacts had appeared. All steps in MeshLab, except the orientation of the tooth into occlusal view-step, were automated by running MeshLab custom scripts written by the first author (see Supplementary Information for a more detailed Slicer, Meshmixer, and Meshlab protocol and for all MeshLab scripts used in this study).

Final surface meshes were exported as .ply files in ASCII format (unticking the ‘binary encoding’ box in MeshLab).

### Dental topographic calculation

Values for the following DTMs were calculated for all surface meshes in the R package molaR (v5.3, Pampush et al. 2016, 2022)curvature measured as Dirichlet Normal Energy (or “DNE”: Bunn et al. 2011), outwardly facing curvature measured as Convex DNE (Pampush et al. 2022), relief measured as the Relief Index (or “RFI”: Boyer 2008), complexity measured as the Orientation Patch Count Rotated (or “OPCR”: Evans et al. 2007; Evans and Jernvall 2009), the average change in elevation measured as the Slope (Ungar and M’Kirera 2003). The 3D and 2D surface areas of each m2 specimen were automatically output as well. We note that ariaDNE is a variation on DNE that is more robust and consistent under different surface mesh acquisition methods and preparation procedures compared to DNE (Shan et al. 2019). However, as ariaDNE is calculated in MATLAB (2021) which is proprietary software requiring a priced licence fee, we refrained from using it in this study. DTM values were calculated using the ‘molaR_Batch’ function with default settings, except setting ‘findAlpha’ to ‘TRUE’ in order to find the optimal alpha value of each surface mesh for RFI calculation (see Supplementary Information for R script). Upon calculating topographic metrics, several specimens returned an error when finding the alpha value for calculating RFI. These surface meshes (*Saimiri boliviensis* AMNH76003 smoothed using the HCL, TAU10, TAU50, and TAU100 smoothing settings, and *Lagothrix lagothrix* USNM545887 using the TAU100 setting) were read into R using the ‘vcgPlyRead’ function of the ‘Rcvg’ R package (v0.22.1, Schlager 2017), and then had their RFI successfully calculated using the ‘RFI’ function of the molaR package (v5.3, Pampush et al. 2016) and setting the ‘alpha’ value to 0.09, 0.1, 0.09, 0.09, and 1.15, respectively, with these values found via trial and error. *Callithrix jacchus* specimen USNM259427 with the TAU100 smoothing setting produced an error regarding counting arcs when attempting to calculate the RFI. This surface mesh was opened in MeshMixer and checked using the ‘Inspector’ tool. Problematic areas were fixed using the ‘smooth fill’ setting, and 2 triangles were manually removed using the ‘Select’ tool. There were no issues on the updated surface mesh.

### Pipeline validation

Classification accuracies were calculated using a quadratic discriminant analysis (which, unlike a linear discriminant analysis, does not require the variances of the dental metrics to be equal) in R using the ‘qda’ function of the MASS R base package (v7.3-56, Ripley et al. 2013), and a jackknife (‘leave-one-out)’ procedure. We validated our pipeline using the four different smoothing settings mentioned above (HCL, TAU10, TAU50, and TAU100), and different combinations of some or all the following variables, following Winchester et al. (2014): DNE, RFI, OPCR, and the natural log of m2 area.

### Dietary classification scheme

#### Organising raw dietary data

As noted by Cooke (2011), platyrrhine diets can differ markedly between seasons, which poses challenges when assigning strict categories to them. To capture the full breadth of platyrrhine diets, including seasonal differences, a broad sample of 98 primary reports on platyrrhine diets, comprising both published papers and unpublished theses, was examined and each study was entered into a GoogleSheet. Studies were identified based on the dietary compilations of Miranda and Passos (2004), Youlatos (2004), Digby et al. (2006), Norconk et al. (2009), Edmonds (2016), and Janiak et al. (2018), and primary publications mentioned in these were examined to extract data directly from those (see Supplementary Information for full list of references). Mostly these data were presented as the percentage of time feeding, which included foraging time in some cases. The wide range of food items and different dietary groupings used in the different publications were initially consolidated into six broad categories: ‘Fruits’, (which included flowers and fungi), ‘Leaves’, ‘Seeds’, ‘Animal matter’, ‘Exudates’, ‘Fungi’, and ‘Other’. When ‘Fungi’ was used as a separate dietary category, the cluster analysis placed *Callimico* apart from all other taxa, with it being the only member of the ‘Fungivore’ category. However, in order to test whether different taxa with similar diets have convergent dental shapes, we needed at least two taxa per dietary group. As there are no other specialised fungivores besides *Callimico* within Platyrrhini, we grouped Fungi with Fruits based on their relatively similar compositional properties (Norconk et al. 2009). See Table S1 and Supplementary Information for more information on categories and raw data.

Five dietary entries out of >650 were excluded from our dataset because they did not clearly correspond to the five categories defined above: “corn from surrounding plantations” (single occurrence); “soil from termitaria nest, fungi, and a frog” (single occurrence); “fluids of unripe palm nuts (*Astrocaryum*)” (single occurrence); “Buds” (or “broto” in Spanish studies) without further specification whether these are leaf buds or flower buds (two occurrences). In each of these cases, the percentage represented by the excluded entry was added to the remaining dietary components in equal proportions.

#### Cluster analysis of diet

As dietary data was relatively similar within genera (see dietary data per species in SM; see also Rosenberger 2020), we averaged these data per genus (see Table S2 in the Supplementary Information for a list of the species that were included for every genus). For this and all later analyses, we used *Saguinus* sensu lato as a taxonomic unit, which includes newly erected genera *Tamarinus* and *Oedipomidas* (Brcko et al. 2022). We excluded the ‘Other’ category from our further analyses. The genus averages of the remaining five dietary components ‘Fruits’, ‘Leaves’, ‘Seeds’, ‘Animal matter’, and ‘Exudates’ were standardised in R using the ‘scale’ function of the base R base package (R v4.2.0), and a Principal Component Analysis was run on the standardised five variables using the ‘prcomp’ function of the ‘stats’ R base package. Following Pineda-Munoz and Alroy (2014), we calculated the Euclidean distance among the five PC scores using the ‘pca2euclid’ function from the tcR R package (v2.3.2, Nazarov et al. 2015). An Unweighted Pair Group Method with Arithmetic Mean analysis (UPGMA) was performed on the Euclidean distance matrix using the ‘upgma’ function of the R-package phangorn (v2.11.1, Schliep 2011) using default settings. This resulted in the UPGMA tree that was used to identify clusters of platyrrhine taxa with similar diets. The raw dietary data were examined to identify what dietary components characterise each cluster, and this was used when naming our dietary categories.

### Analysis and visualisation of dental topographic metrics

When analysing the entire platyrrhine sample, we excluded five moderately to heavily worn non-callitrichid specimens of the Winchester et al. (2014) sample. These five specimens were: *Ateles belzebuth* USNM241384, *Cacajao calvus* AMNH98316, *Cebus capucinus* USNM291133, *Lagothrix lagotricha* AMNH71767, and *Lagothrix lagotricha* USNM545878.

One-way ANOVAs were run using the ‘aov’ function of the R base package (R v4.2.0, R Core Team 2022). When a significant difference was found between categories, a Games-Howell post hoc test to account for unequal sample sizes and variances was applied using the ‘games_howell_test’ function of the rstatix package (v0.7.2, Kassambara 2023). A Principal Component Analysis (PCA) was run using the following variables: Convex DNE, OPCR, RFI, Slope, and the natural log of the 2D crown area and using the ‘prcomp’ function of the stats package (R v4.2.0, R Core Team 2022), setting scale to ‘TRUE’ to standardise all variables before conducting the PCA.

The entire sample of platyrrhine surface meshes (excluding the heavily worn specimens mentioned above) was analysed to compare dental topography of callitrichids to non-callitrichid platyrrhines, and to test the accuracy of dietary prediction when callitrichids were added. First, we analysed the entire platyrrhine sample for the four differently smoothed datasets to determine which smoothing setting yielded the highest classification accuracy using the following five variables individually, and also the combination of these: DNE or Convex DNE, RFI, OPCR, Slope, and the natural log of 2D m2 area. The smoothing setting that yielded the highest classification accuracy was identified and used for the further analyses and descriptions of callitrichid dental topography. For the final quadratic discriminant analyses for predicting diet on the entire sample, we used topographic variables only (DNE or Convex DNE, RFI, OPCR, and Slope), and compared these results to including topographic variables plus the natural log of 2D m2 area. These final analyses were performed using three different dietary classification schemes: our original UPGMA scheme, an edited-UPGMA scheme (see Results below), and the scheme used by Winchester et al. (2014).

## Results

### Pipeline validation results using non-callitrichid platyrrhine sample and Winchester et al. (2014) dietary classification scheme

A comparison of the classification accuracy of quadratic discriminant analysis (QDA) performed on surface meshes derived from our new freeware dental topography pipeline compared to those of Winchester et al. (2014, table 7) is shown in Table 2. When topographic variables were analysed separately, a stark difference appears in the classification accuracies of DNE and OPCR between the different processing protocols. Results from Winchester et al. (2014) show a relatively high classification accuracy using DNE only (67.7%), but relatively low for OPCR only (44.1%). In contrast, our pipeline results in DNE having a relatively low classification accuracy (ranging from 37.84% to 47.75%, depending on the smoothing setting used) and varied classification accuracy of OPCR (ranging from a relatively high accuracy of 68.47% to a low accuracy of 29.73%). RFI yields relatively high classification accuracies across all protocols.

**Table 2.**
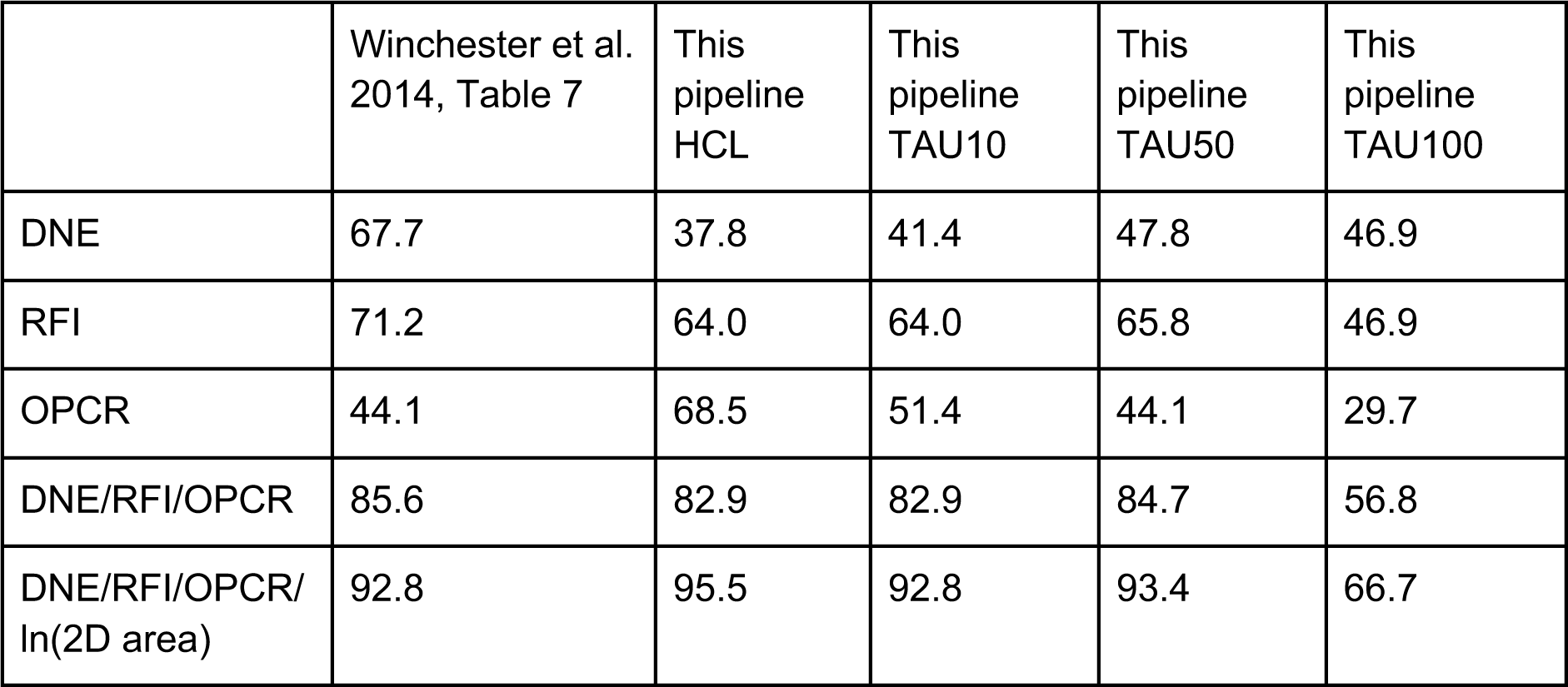
Pipeline validation: classification accuracy of non-callitrichid platyrrhine sample (n=111). This is the original sample of Winchester et al (2014, morphosource.org project ID: 000000C89), and so includes the five specimens that were excluded from the remainder of this study (due to excessive wear). Dietary groups followed Winchester et al. (2014)

When topographic values were combined (DNE, RFI, and OPCR), our pipeline performed comparably well, albeit slightly worse, than that of Winchester et al. (2014, 85.6%), but only by <3% for the HCL (82.88%), TAU10 (82.88%), and TAU50 (84.68%) smoothing settings. The three topographic variables combined using our pipeline with the TAU100 smoothing setting results in a considerably lower classification accuracy of 56.76%. When the natural log of m2 length is included (following Winchester et al. 2014), all versions of the pipeline reach their highest classification accuracies. Except when using the TAU100 smoothing setting, classification accuracies of the three topographic variables and a measure of size were comparable for all processing pipelines, with our pipeline using the HCL smoothing setting outperforming Winchester et al. (2014)’s protocol, albeit by less than 1% (93.69% versus 92.8%, respectively, see Table 2).

### Dietary classification scheme

The results of the UPGMA analysis of dietary information extracted from 98 studies are shown in Figure 2. Some callitrichid taxa (*Cebuella* and *Mico*) were not included in the UPGMA analysis as they were missing from the compilations we consulted, but were included a posteriori, as discussed below. Based on the UPGMA tree, we identify five primary dietary clusters within Platyrrhini: Frugivory-Insectivory (*Saimiri, Callimico, Cebus, Leontocebus, Saguinus* sensu lato*, Leontopithecus*), Frugivory (*Plecturocebus, Aotus, Pithecia, Callicebus, Ateles*), Seed eating (*Chiropotes, Cheracebus, Cacajao*), Folivory (*Brachyteles, Alouatta*), and Exudate feeding (*Callithrix*). The dietary data for each of these genera and clusters are shown in Table 3. The clusters can be characterised as follows: Frugivore-insectivores have a large component of fruits in their diet (≥44%) and also a considerable component of insects in their diet (≥24%, except for *Leontocebus*, which has an average insect intake of 13%) and low intake of leaves for most genera (≤6%, except for *Sapajus* with 25%). The frugivores are characterised by having over 58% of their diet consisting of fruits, with the second largest dietary component comprising a moderately high intake of leaves (12-28%), and their insect intake is low (<12%). The folivores have diets comprising >52% leaves, and a large secondary component of fruit in their diet (40-46%). Seed eaters have a diet that consists of >32% seeds, and the diet of exudate feeders consists of 52% of exudates. This dietary classification scheme is referred to as ‘UPGMA-based’.

**Fig. 2.**
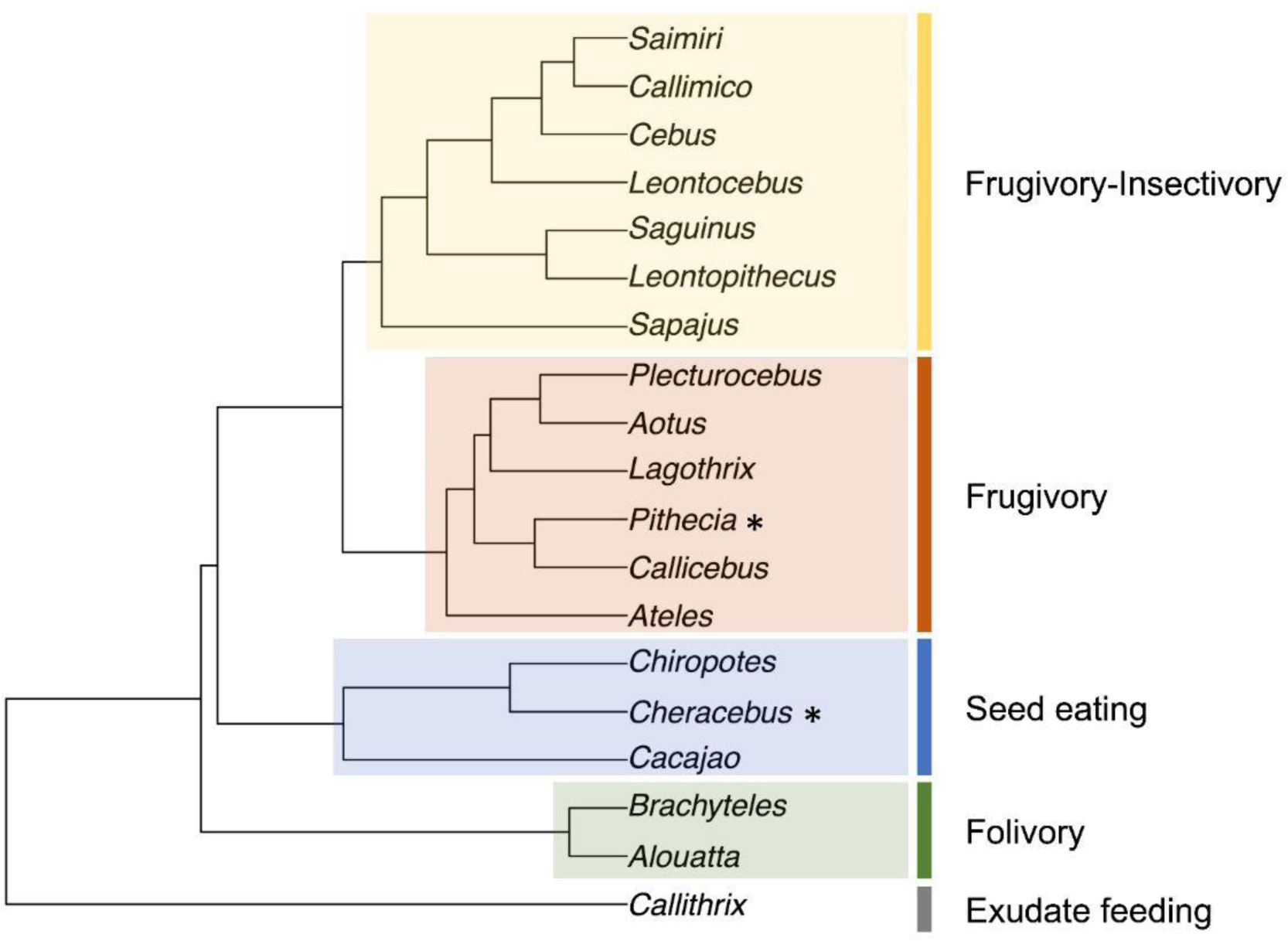
UPGMA results with an added colour scheme to highlight the clusters we identified as dietary categories. *Pithecia* and *Cheracebus* are marked with an asterisk as their dietary category was altered in the edited-UPGMA dietary scheme in which *Pithecia* was classified as a seed feeder and *Cheracebus* as a frugivore

**Table 3.** Genus averages of dietary data used for UPGMA (note that the ‘Other’ category was not included in the UPGMA and that the categories ‘Fruits’, ‘Flowers’, and ‘Fungi’ were combined into a single ‘Fruits’ category; see Table S1 for the breakdown of diets including these categories). Raw input data of each genus are grouped per dietary cluster in the left-hand column based on the UPGMA output. *Cheracebus* and *Pithecia* are marked with an * as they were placed in another group for the edited UPGMA scheme

| Genus | Fruits | Leaves | Seeds | Animal matter | Exudates | Other |
| --- | --- | --- | --- | --- | --- | --- |
| **Exudate feeding** |  |  |  |  |  |  |
| *Callithrix* | 0.28 | 0 | 0 | 0.21 | 0.52 | <0.01 |
| **Seed eating** |  |  |  |  |  |  |
| *Cacajao* | 0.19 | 0.04 | 0.69 | 0.04 | 0 | 0 |
| *Cheracebus* * | 0.47 | 0.09 | 0.32 | 0.09 | 0 | 0 |
| *Chiropotes* | 0.45 | 0.04 | 0.45 | 0.02 | 0 | 0.04 |
| **Folivore** |  |  |  |  |  |  |
| *Alouatta* | 0.40 | 0.57 | <0.01 | <0.01 | 0 | 0.01 |
| *Brachyteles* | 0.46 | 0.52 | 0 | 0 | 0 | 0 |
| **Frugivore** |  |  |  |  |  |  |
| *Ateles* | 0.83 | 0.12 | 0.02 | <0.01 | 0 | 0.02 |
| *Callicebus* | 0.70 | 0.16 | 0.13 | 0 | 0 | <0.01 |
| *Aotus* | 0.64 | 0.24 | 0 | 0.03 | 0 | 0 |
| *Lagothrix* | 0.69 | 0.12 | 0.05 | 0.12 | 0 | 0.01 |
| *Plecturocebus* | 0.58 | 0.28 | 0 | 0.10 | 0 | <0.01 |
| *Pithecia* * | 0.60 | 0.17 | 0.21 | 0.02 | 0.01 | <0.01 |
| **Frugivore-Insectivore** |  |  |  |  |  |  |
| *Callimico* | 0.58 | 0 | 0 | 0.34 | 0.01 | 0.07 |
| *Cebus* | 0.58 | 0.06 | 0.03 | 0.27 | 0 | 0.03 |
| *Leontocebus* | 0.56 | 0 | 0 | 0.27 | 0.12 | 0.01 |
| *Leontopithecus* | 0.79 | 0 | 0 | 0.13 | 0.08 | <0.01 |
| *Saguinus* | 0.65 | 0 | 0 | 0.24 | 0.07 | 0.02 |
| *Saimiri* | 0.64 | 0 | 0.01 | 0.31 | 0 | 0 |
| *Sapajus* | 0.44 | 0.25 | 0.05 | 0.27 | 0 | <0.01 |

The placements of *Cheracebus* and *Pithecia* in the UPGMA tree shown in Figure 2 are somewhat surprising given they differ from other dietary schemes (Cooke 2011; Winchester et al. 2014; Allen et al. 2015). In our results, *Cheracebus* is grouped with the seed eaters, separate from the other callicebine genera *Callicebus* and *Plecturocebus*, which are grouped in the frugivore cluster. *Pithecia* is grouped among frugivores, in contrast with the other pitheciine genera *Cacajao* and *Chiropotes*, which are grouped as seed eaters instead. Our grouping may be driven by the low number of observational studies in our consulted literature (*Cheracebus,* n = 2), data used from possibly unrepresentative habitats and thus possibly unrepresentative dietary data, as well as seeds being grouped with fruits in several studies but not in ours (*Pithecia*). We therefore also used an alternative dietary classification scheme (referred to as edited-UPGMA) in which *Cheracebus* is classified as a frugivore and *Pithecia* as a seed eater (see Table 1 and 4), congruent with field studies regarding their diet (e.g., Peres 1993; Norconk 1996 that are included in our UPGMA but are only two out of 11 *Pithecia* reports used in our study). We considered the edited-UPGMA scheme as our preferred dietary scheme for subsequent analyses.

The callitrichid taxa (*Cebuella* and *Mico*) that were not included in the UPGMA analysis but are present in the dental topographic sample, namely *Cebuella* and *Mico*, were classified as exudate feeders (and Tavares 1999; Veracini 2009 for *Mico*; following Kay et al. 2019 for *Cebuella*). As the callitrichid taxa were not included by Winchester et al. (2014), they were therefore not assigned dietary categories in their original classification scheme. For the final quadratic discriminant analyses on the entire platyrrhine sample, callitrichids in the ‘diet Winchester’-scheme were assigned to the same groups as in the UPGMA dietary schemes (both the UPGMA-based and the edited-UPGMA scheme are the same regarding the callitrichid categories), or the closest group used in the original Winchester et al. (2014) classification scheme. Thus, for these final analyses, all exudate feeders are classified as part of the exudate feeding-group (a dietary category originally not present in Winchester et al. 2014), and frugivore-insectivore callitrichids are classified as part of the insectivore-omnivore-group in ‘diet Winchester’. *Sapajus* was not considered as a separate genus from *Cebus* by Winchester et al. (2014), and so we assign it the same dietary category as *Cebus* in their scheme (i.e., Hard-Object feeding). See Table 4 for an overview of the dietary categories used for each genus in each of the three classification schemes.

**Table 4.** Dietary classification schemes: ‘UPGMA-based’ and ‘edited-UPGMA’ use dietary grouping resulting from the UPGMA (see text), ‘Winchester et al.’ is following Winchester et al. (2014, Table 1). Callitrichids that were added to the sample were assigned dietary groups following the UPGMA or the closest equivalent of the Winchester et al. dietary classification scheme. Categories in bold indicate the differences between the UPGMA-based and the edited-UPGMA schemes

| **Genus** | **UPGMA-based** | **edited-UPGMA** | **Winchester et al.** |
| --- | --- | --- | --- |
| *Alouatta* | Folivore | Folivore | Folivore |
| *Aotus* | Frugivore | Frugivore | Frugivore |
| *Ateles* | Frugivore | Frugivore | Frugivore |
| *Brachyteles* | Folivore | Folivore | Folivore |
| *Cacajao* | Seed eating | Seed eating | Hard-Object feeding |
| *Callimico* | Frugivore-Insectivore | Frugivore-Insectivore | Insectivore-Omnivore |
| *Callithrix* | Exudate feeding | Exudate feeding | Exudate feeding |
| *Cebuella* | Exudate feeding | Exudate feeding | Exudate feeding |
| *Cebus* | Frugivore-Insectivore | Frugivore-Insectivore | Hard-Object feeding |
| *Cheracebus* | **Seed eating** | **Frugivore** | Frugivore |
| *Chiropotes* | Seed eating | Seed eating | Hard-Object feeding |
| *Lagothrix* | Frugivore | Frugivore | Frugivore |
| *Leontocebus* | Frugivore-Insectivore | Frugivore-Insectivore | Insectivore-Omnivore |
| *Leontopithecus* | Frugivore-Insectivore | Frugivore-Insectivore | Insectivore-Omnivore |
| *Mico* | Exudate feeding | Exudate feeding | Exudate feeding |
| *Pithecia* | **Frugivore** | **Seed eating** | Hard-Object feeding |
| *Plecturocebus* | Frugivore | Frugivore | Frugivore |
| *Saguinus* | Frugivore-Insectivore | Frugivore-Insectivore | Insectivore-Omnivore |
| *Saimiri* | Frugivore-Insectivore | Frugivore-Insectivore | Insectivore-Omnivore |
| *Sapajus* | Frugivore-Insectivore | Frugivore-Insectivore | Hard-Object feeding |

Our preferred edited-UPGMA dietary scheme differs little from that of Winchester et al. (2014); their hard-object feeder category is broadly equivalent to our seed eating category (except for *Cebus* and *Sapajus*, discussed below), and their insectivore-omnivore category is directly equivalent to our frugivore-insectivore category. The only differences between our edited-UPGMA scheme and that of Winchester et al. (2014) are the classifications of *Cebus* and *Sapajus* as hard-object feeders by Winchester et al. (2014) and as frugivore-insectivores in our edited-UPGMA scheme. Based on the studies used in our dietary classification scheme, *Cebus* and *Sapajus* have a low seed intake (3 and 5%, respectively) and thus are not grouped with other seed eaters in the UPGMA (which had seed intakes of 32-69%, see Table 3). Our classification of *Cebus* and *Sapajus* as frugivore-insectivores is driven by their moderate fruit intake with a considerable secondary component of insects (see Table 3). We did not take the mechanical properties, such as ‘hardness’ or ‘toughness’, of food into account in our scheme, unlike Winchester et al. (2014) who accommodate this in their ‘hard-object feeder’ category, and they placed *Cebus* and *Sapajus* in this group based on the highly mechanically challenging materials in their diet, such as fruits with tough exocarp. We note that some other published platyrrhine dietary classification schemes have placed *Cebus* and *Sapajus* separately from the seed-eating *Pithecia*, *Cacajao,* and *Chiropotes*, being instead classified as having a ‘frugivore/omnivore’ (Cooke 2011), ‘frugivore’ (Allen et al. 2015) or ‘frugivore/seed’ (Ungar et al. 2018) diet.

### Dental topography of callitrichids

Raw image stacks of the scanning data and surface meshes cropped at the base of the crown along the cementum-enamel junction of the 39 callitrichid m2 specimens are now publicly available on MorphoSource (project ID:000471738). The fully processed surface meshes of the entire platyrrhine sample (n=150) using our pipeline and the HCL smoothing setting (which resulted in the highest classification accuracy, see above) are now also publicly available on MorphoSource (project ID: 000471738).

When analysing the entire platyrrhine sample (excluding five moderately worn specimens that are part of the original Winchester et al. 2014 sample and including 39 new callitrichid specimens) using our pipeline with four different smoothing settings, the HCL smoothing setting resulted in the highest classification accuracy, consistently outperforming all three TAU smoothing settings by 4 to 12% (see Table 5). The different iteration settings of the Taubin smooth setting show little difference in classification accuracy between 10 and 50 iterations for every variable, but show a great decrease in accuracy for 100 iterations for every variable except DNE. Differences between using DNE (including both convex and concave DNE) versus Convex DNE (which only includes the convex DNE and has been argued to reflect a functional signal better as it only takes outwardly facing curvature into account, Pampush et al. 2022) on this sample are minimal, with the largest difference being 3.45% (see Table 5).

**Table 5.**
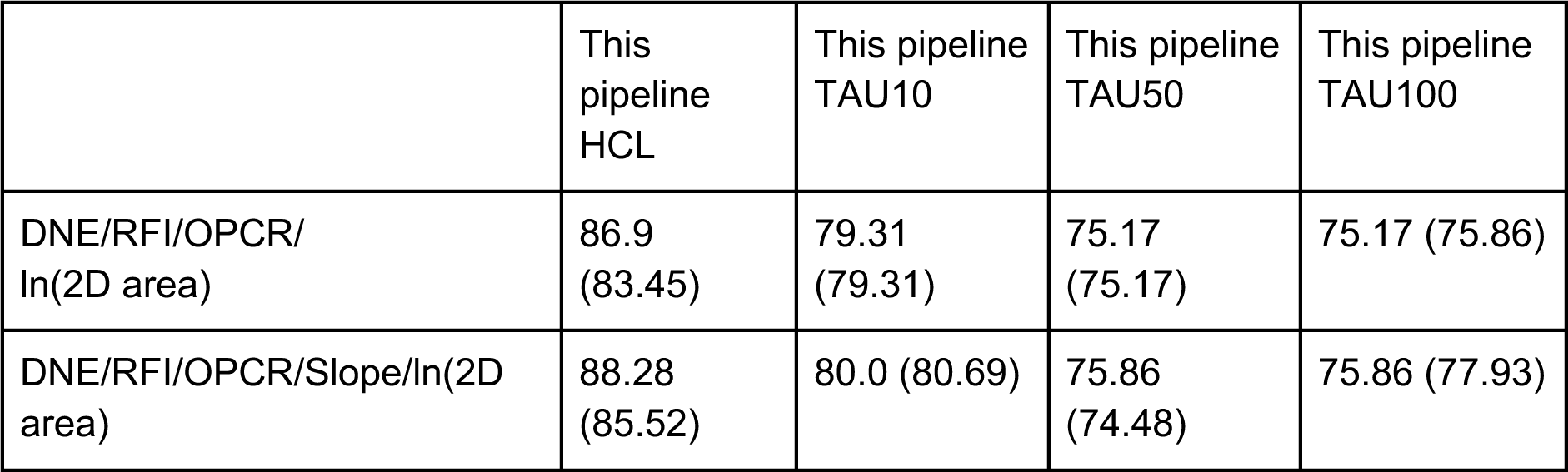
Classification accuracy percentages per different smoothing settings for a combination of DTMs; the QDA was run on the entire sample (n-total=145, callitrichids included, five non-callitrichid specimens excluded) using the edited-UPGMA dietary categories. The values in parentheses are the classification accuracy percentages of the same variables, except that instead of convex DNE, DNE of the entire crown (i.e., both convex and concave) is used

Figure 3 shows each topographic metric, as well as size as a boxplot and violin plot per dietary category following the edited UPGMA scheme. One-way ANOVAs indicate a significant difference between the dietary categories for ln(2D area), OPCR, RFI, and Slope, whereas DNE and Convex DNE do not differ significantly between dietary categories (see Table S3 for ANOVA results). Post hoc Games-Howell tests were applied to the variables that differed significantly between dietary categories, and all significantly different pairs are marked in Figure 3 (see Table S4 for all Games-Howell post hoc results). The significantly different variables between diets are discussed below.

**Fig. 3.**
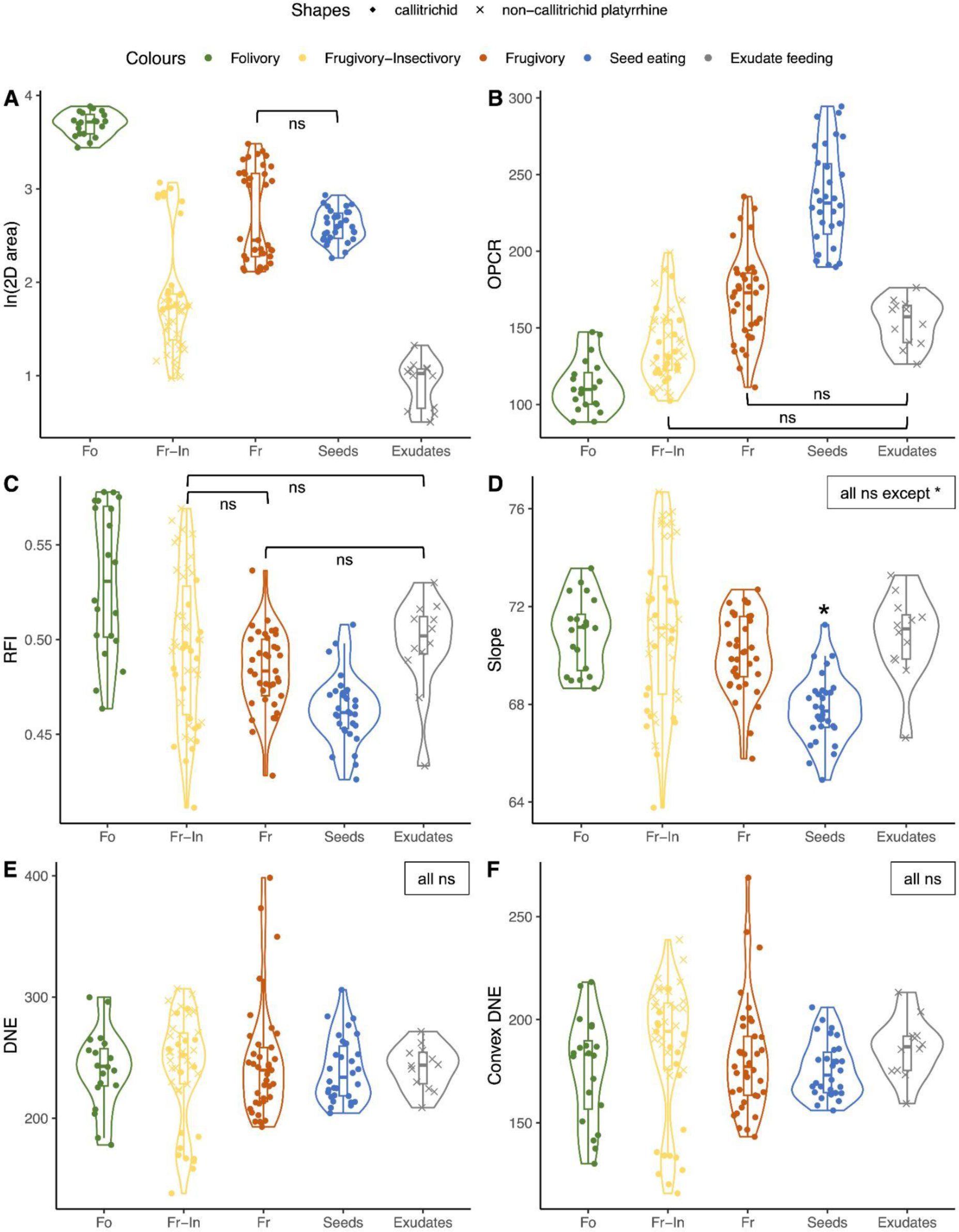
Boxplots of a) the natural log of (2D area); b) OPCR, c) RFI, d) Slope, e) DNE, and f) Convex DNE per different dietary category following the edited-UPGMA scheme (see text). Fo = folivory; Fi-In = Frugivory-Insectivory; Fr = Frugivory. Pairwise comparisons were significant (p < 0.05) in Games-Howell post hoc analyses except for those indicated as not significant (‘ns’). The metric Slope (d) only differed significantly between the Seed eating category and all other dietary categories

Size, measured as the natural log of the 2D crown area, differs significantly between all dietary categories except between frugivores and seed eaters (see Figure 3a). Although significantly different between categories, there is overlap in size between the largest frugivore-insectivores and the frugivores and seed eaters (Figure 3a, see Table S5 for the range of each DTM per dietary group). This overlap in size is solely due to the relatively large *Cebus* and *Sapajus* specimens in the frugivore-insectivore category. The newly added callitrichids represent the smallest members of the frugivore-insectivore category, and expand its size range downwards.

OPCR differs significantly between dietary categories, except between exudate feeders and frugivores, and between exudate feeders and frugivore-insectivores. Following results of Winchester et al. (2014), folivores have the lowest OPCR values, followed by the frugivore-insectivores. Frugivores have low to medium OPCR values, and seed eaters have the highest complexity values (see Table S5). Exudate feeders are characterised by intermediate values for OPCR, overlapping with frugivores and frugivore-insectivores (see Figure 3b, Table S5). The newly added frugivore-insectivore callitrichids are found throughout the range of OPCR values of other (non-callitrichid) frugivore-insectivores.

RFI shows large ranges for, and considerable overlap between, each dietary category (see Figure 3c, Table S5). RFI only differs significantly between folivores and all other categories (folivores having higher RFI), and between seed eaters and all other categories (seed eaters having lower RFI). Frugivore-insectivores are characterised by the largest range in RFI values, almost covering the entire RFI range of the total sample. The newly added frugivore-insectivore callitrichids are present throughout the entire range of other, non-callitrichid frugivore-insectivores and expand the category’s range upwards due to the high RFI values of *Callimico*.

All diets overlap in values of Slope, except for seed eaters, which differ from all other dietary categories by having significantly lower Slope (see Figure 3d, Table S5). Frugivore-insectivores show the largest range in Slope values, with the newly added callitrichids expanding the range upwards: the highest Slope of the entire sample is mostly driven by specimens of *Callimico*, some *Saguinus* specimens and a single *Leontocebus* specimen.

Our results allow us to characterise the dental topography of exudate-feeding platyrrhines (*Callithrix, Cebuella,* and *Mico*). Compared to other extant platyrrhines, the m2s of exudate feeders are characterised by a combination of small size (range: 0.51-1.33), a medium-low OPCR (range: 126-176), and medium-high RFI (range: 0.43-0.53).

### Callitrichid dental topography compared to that of other platyrrhines

The PCA plot of the variables Convex DNE, OPCR, RFI, Slope, and the natural log of the 2D crown area is shown for PC1 and PC2 (together capturing 77.61% of total variation) in Figure 4. This plot shows variable degrees of overlap or separation between the different dietary categories. PC1 correlates positively with OPCR (0.43), and negatively with RFI (-0.59) and Slope (-0.61). PC2 correlates positively with size (0.64) and negatively with Convex DNE (-0.62, see Figure 4). All folivores occupy a space that reflects their larger m2 size at the top half of the graph (higher PC2 values) and in general a lower OPCR (lower PC1 values) than the other specimens. The seed eating category shows a combination of higher OPCR and lower RFI values (higher PC1 values) compared to other categories. Frugivores and frugivore-insectivores occupy a large area covering the middle of the PCA plot. The frugivores are split into two clusters based on size (see also Figure 3a), with the large Frugivores *Ateles* and *Lagothrix* overlapping with part of the folivore cluster, and the small frugivores *Cheracebus*, *Plectorucebus*, and *Aotus* occupying the middle of the PCA plot, overlapping with some of the seed eaters, frugivore-insectivores, and exudate feeders. The frugivore-insectivore group can be roughly split into three clusters: 1) in the top right space of the PCA plot, a cluster solely consisting of *Cebus* and *Sapajus* is found, driven by their large size; 2) in the middle of the PCA plot, overlapping with exudate feeders and the small bodied frugivores, occupied by numerous specimens of callitrichid *Leontopithecus* and *Saguinus*, one *Callimico* specimen, and the non-callitrichid platyrrhine *Saimiri*, driven by their small size and medium OPCR; 3) on the left of the PCA plot, a cluster of frugivore-insectivores is formed by callitrichids *Callimico* and *Saguinus*, driven by the combination of their small size and exceptionally low RFI. Exudate feeders overlap with the second, or middle, cluster of frugivore-insectivores due to their small 2D crown areas in combination with medium values of OPCR, Convex DNE, and RFI. Finally, seed eaters cluster on the right side of the PCA plot, driven by high OPCR values and low values of RFI and Slope.

**Fig. 4.**
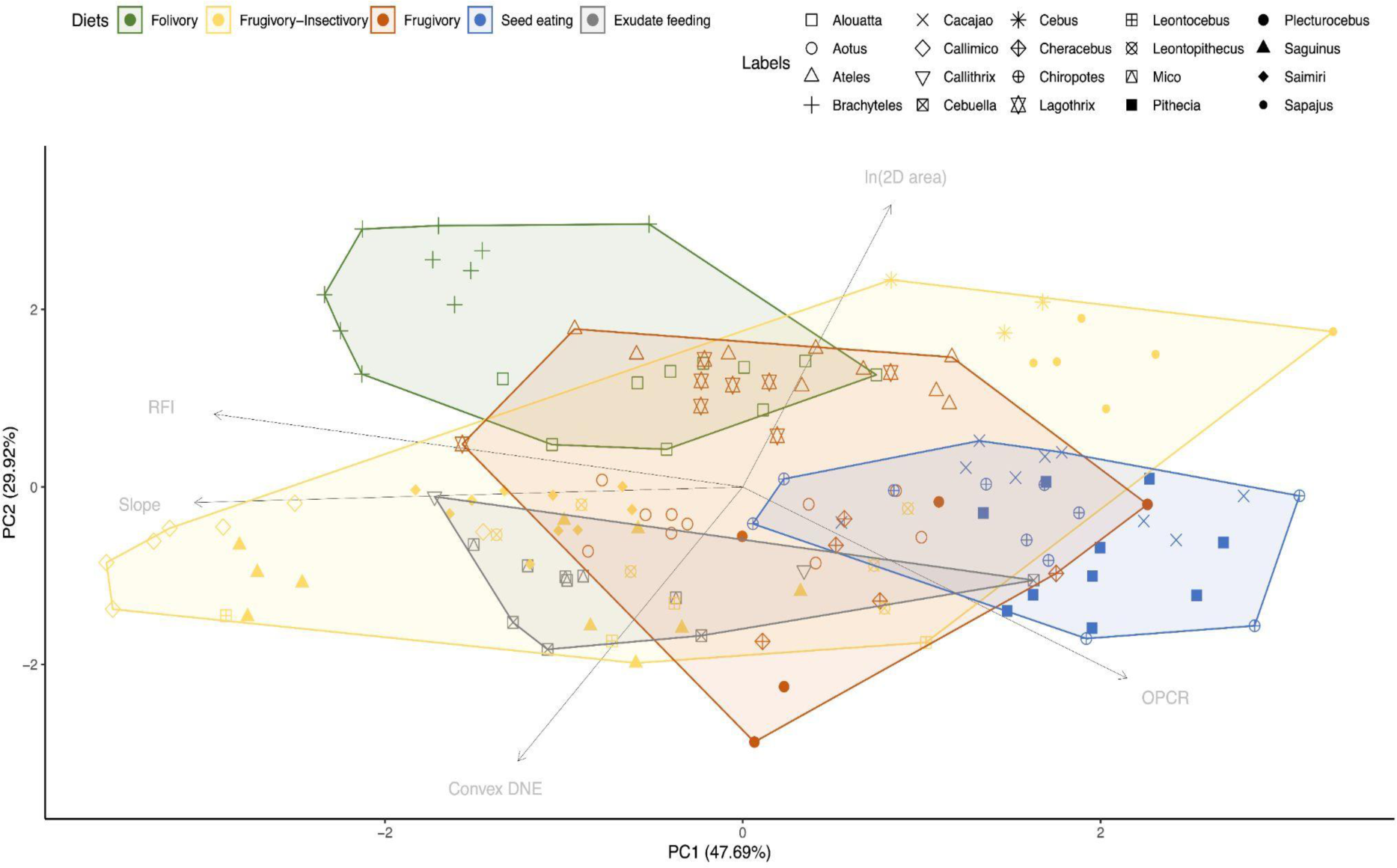
PCA of variables Convex DNE, OPCR, RFI, Slope and the natural log of the 2D crown area. PC1 (47.69%) and PC2 (29.92%) capture 77.61% of the total variance. The PC loadings are plotted with an arrow for each variable in light grey

### Dietary classification accuracies using dental topography of platyrrhines

The edited-UPGMA dietary scheme classification consistently out-performed the original UPGMA scheme in terms of classification accuracy by more than ten percent (13.10-17.09%, see Table 6). The edited-UPGMA scheme underperformed slightly compared to the success of the classification scheme of Winchester et al. (2014), with the largest difference in performance being 8.49% (see Table 6). This maximum difference decreased when a measure of molar size was included, and the edited-UPGMA classification scheme performed equally well as that of Winchester et al. (2014), or underperformed by a maximum of 4.48%.

**Table 6.** Classification accuracy percentages of meshes processed using the new pipeline (n = 145). “Topography” = Convex DNE, RFI, OPCR, and Slope; “Size” = natural log of 2D area. The values in parentheses are the classification accuracy percentages of the same variables, except that instead of convex DNE, DNE of the entire crown (i.e., both convex and concave) is used

| HCL smoothing | UPGMA-based | Edited-UPGMA | Winchester et al. |
| --- | --- | --- | --- |
| **Platyrrhines without callitrichids (n=106)** |  |  |  |
| **Topography** | 64.15 (66.04) | 81.13 (81.13) | 88.68 (89.62) |
| **Topography + size** | 80.19 (78.30) | 95.28* (92.45) | 95.28 (94.34) |
| **Platyrrhines with callitrichids (n=145, including extra dietary category ‘Exudate feeding’)** |  |  |  |
| **Topography** | 60.0 (61.38) | 73.10 (73.79) | 77.93 (79.31) |
| **Topography + size** | 80.69 (79.31) | 90.34** (90.34) | 94.48 (93.10) |

\*Break-down of classification accuracy per dietary category of this test: Folivory = 95%, Frugivore-Insectivore = 89.47%, Frugivory = 100%, and Seed eating = 93.33%.
\*\*Break-down of classification accuracy per dietary category of this test: Folivory = 95%, Frugivore-Insectivore = 84.78%, Frugivory = 97.30%, Seed eating = 93.33%, and Exudate feeding = 75%.

When the callitrichid specimens were added to the sample, and an additional category (exudate feeding) was added to the dietary classification scheme, classification accuracy drops in nearly all cases by 1% to 11% (Table 6), with the only exception being topography + size for the UPGMA dietary scheme. However, overall the classification accuracy is good, ranging from 80 to 95% when a measure of size is included. The decrease in classification accuracy is mainly driven by the misclassification of exudate feeders (which has the lowest classification accuracy of any dietary category, namely 75%), and reduced classification accuracy of frugivore-insectivores (by 4.7%) and frugivores (by 2.7%, see Table S6 and S7). When considering the classification results of individual specimens (see Supplementary Information), it becomes clear that the increased misclassifications are mostly driven by the increased confusion of frugivore-insectivores with other categories.

## Discussion

### Freeware pipeline validation

Our results indicate that the freeware pipeline presented here produces similar classification accuracies to those produced by the protocol of Winchester et al. (2014) when combining DTMs. As our freeware pipeline replicates results from proprietary software, the protocol recommendations provided by Spradley et al. (2017), Berthaume et al. (2019b), and Melstrom and Wistort (2021) regarding the effects of smoothing, cropping, and simplification still apply to our freeware pipeline as well.

However, when considering the topographic variables in isolation for dietary classification, our pipeline produces noticeably different results from those of the protocol of Winchester et al. (2014) for DNE and OPCR values. Specifically, the results of Winchester et al. (2014) show a relatively high classification accuracy for DNE by itself, and a low classification accuracy for OPCR by itself. Our results support the opposite; OPCR separates the dietary categories relatively well, whereas DNE does not (see Results and Figure 3e, b, and f). We suspect that this is due to the different smoothing functions applied in MeshLab compared to that in Amira (or Avizo), which in our proposed pipeline presumably removes some informative bending energy information (i.e., DNE) and thus reduces the signal of functional differences between the different dietary categories. In contrast, our pipeline picks up on (or retains) features in dental complexity that seem to be missed (or removed) by the proprietary protocol executed in Amira (or Avizo) and GeoMagic Studio, as our results show a stronger dietary signal in the molar complexities. The difference in classification accuracies of OPCR and DNE between our proposed pipeline and that of protocols using proprietary software point at a fundamental difference in the processing of digital surface meshes resulting in different shape characteristics that are quantified by the dental topographic variables. This may make direct comparison between results obtained by using this freeware pipeline and those of proprietary protocols difficult. Morley and Berthaume (2023) also identify this downside when using Meshlab’s smoothing option (Laplacian Smooth) and instead recommend using the vcgSmooth (Taubin) tool within the R package Rvcg. Ultimately, we conclude that our proposed pipeline performs equally well as previous protocols using proprietary software, as the overall classification accuracy when using multiple dental topographic metrics (as would normally be the case) was comparable between the different protocols.

The different iteration settings of the Taubin smooth setting showed that smoothing for 100 iterations reduces the classification accuracy dramatically compared to that of 50 iterations for every variable, except for DNE. This suggests that, whereas smoothing for this many iterations removes functionally informative information of RFI and OPCR, it instead increases the functional signal of DNE. It may be that the samples of the lighter smoothing settings included irregularities in bending energy that are functionally insignificant and potentially misleading, and are reducing the ability of DNE to capture dietary adaptations. Our results suggest a much stronger smoothing (TAU100) results in higher classification accuracy of DNE, although this may be sample specific and other samples need to be tested to confirm whether this is inherent to our protocol. We recommend choosing the smoothing setting out of the different MeshLab smoothing settings based on the sample and research question under study; we emphasise that we do not necessarily think that the HCL smoothing setting is automatically the best setting for every study.

Although our freeware pipeline shares steps with that of the proposed freeware workflow introduced recently by Morley and Berthaume (2023), there are some differences. We include steps regarding surface mesh deformity reconstruction in MeshMixer and orienting the surface mesh into occlusal view, steps that are not included in the protocol by Morley and Berthaume (2023) but are important for dealing with specimens not previously processed for analyses using DTMs. Morley and Berthaume (2023) compared smoothing settings of various freeware packages, but only one smoothing setting in MeshLab, whereas we compare four different smoothing settings within MeshLab only. Finally, as discussed above, our validation study is designed to simply test whether our freeware pipeline produces surface meshes capable of accurately distinguishing between different diets using DTMs, whereas Morley and Berthaume (2023) focussed on identifying a pipeline that is capable of replicating as closely as possible the specific decimation and smoothing steps implemented by Avizo.

### New dietary classification scheme

The preferred edited-UPGMA dietary scheme differs little from that of Winchester et al. (2014, see Results) in terms of which taxa are referred to which dietary category. When the entire sample is considered (n = 145), our edited UPGMA scheme performs slightly worse (by 0-8%, see Table 6) compared to the dietary groupings of Winchester et al. (2014). The only differences in schemes is the classification of *Cebus* and *Sapajus* as a frugivore-insectivore in the edited-UPGMA scheme, but as hard-object feeders by Winchester et al. (2014). However, the increase in misclassifications is only partly driven by the additional misclassification of three *Cebus* and *Sapajus* specimens (one *Sapajus* as a seed eater, and one *Sapajus* and one *Cebus* as frugivores), that are correctly classified using the Winchester scheme. An additional pair of misclassified specimens using the edited-UPGMA scheme, but that are classified correctly in the Winchester scheme, are two *Saguinus* sensu lato specimens that are misclassified as exudate feeders rather than as their assigned category frugivore-insectivores. Our results thus show that by including *Cebus* and *Sapajus* as frugivore-insectivores, it is harder for QDA to correctly classify ‘frugivore-insectivores’. In contrast, when *Cebus* and *Sapajus* are considered seed eaters or ‘hard-object’ feeders, as in the Winchester et al. (2014) scheme, these five specimens are correctly classified into their dietary groups (as hard-object feeders in the case of *Cebus* and *Sapajus*, and as omnivores in the case of *Saguinus* sensu lato). As *Sapajus* in particular exhibits craniofacial adaptations for hard-object feeding (Daegling 1992; Wright 2005), it is perhaps not that surprising this taxon is not being grouped with the other frugivore-insectivores in the QDA.

### Callitrichid dental topography

QDA results are discussed for when using the edited-UPGMA scheme and including both topography and size (see the * marked entries in Table 6). Classification accuracies per dietary category changed only slightly when callitrichids were added to the sample.

The relatively frequent misclassification of exudate feeders as frugivore-insectivores (and vice-versa) is supported by the overlap in molar topography and size of exudate feeder and frugivore-insectivore specimens (shown in Figure 3 and Table S5). For each dental topographic metric, the range of exudate feeding specimens falls completely within the range exhibited by frugivore-insectivores, and in all cases the exudate range is narrower than that of frugivore-insectivores. It is only in m2 size that the exudate-feeding specimens are distinguished and in which their range extends below that of frugivore-insectivores (although there is overlap between the largest exudate feeders and smallest frugivore-insectivores; see Figure 3a and Table S5). This has been noted by Kirk and Simons (2001) to probably be due to primates specialising on exudativory, similar to insectivory, being unable to sustain large body sizes due to being available only in small feeding quantities and being quite limited in the amount that can be harvested per day. We note that dental topography, however, does aid in the successful classification between these groups in some cases, as it is not necessarily the specimens within this range of size-overlap that are the misclassified specimens. For example, the smallest frugivore-insectivore in our sample (*Saguinus midas* specimen USNM393810) is well within the range of exudate feeders, but is classified correctly as a frugivore-insectivore.

Even though the classification accuracy of exudate feeders was the lowest of all dietary categories in our sample (75% accuracy), this is still much higher than chance (one out of five, or 20%) and demonstrates that the molars of the three exudate feeding genera in our sample (*Callithrix, Cebuella, Mico*) show a consistent combination of dental metrics (medium-low OPCR and medium-high RFI, in combination with small size, as measured by 2D area). However, our results also indicate that the teeth of exudate feeders closely resemble those of frugivore-insectivores, and that dental metrics of exudate feeders fall entirely within the range of frugivore-insectivores in all but one metric (size). Our results thus suggest that there are no particularly distinctive topographic adaptations to exudate feeding present in m2s (congruent with the discussion of Fulwood et al. 2021 in a strepsirrhine sample). This is not completely unexpected, since exudates do not require much masticatory processing by the molars, and their physical consistencies are likened to those of extremely soft fruits (Kay and Covert 1984). The reduction-to-loss of last molars is proposed as a mammalian dental signature for exudate-feeding by Burrows et al. (2020). All callitrichids have lost their final molars (m3s) except *Callimico,* although not all callitrichids are obligate exudate feeders and frugivore-insectivore callitrichids also lack an m3. Instead, adaptations for an exudate and insectivorous diet are located in the anterior dentition, such as procumbent lower incisors with sharp ‘gouging’ edges due to the lack of lingual enamel on incisors (Wible and Burrows 2016; Rosenberger 1978; Francisco et al. 2017; Burrows et al. 2020), and significantly larger incisors and canines compared to the molar sizes (Natori and Shigehara 1992); all of these features facilitate gouging and removing tree bark to stimulate exudate flow, but also to access insects (Rosenberger 1992). As exudate feeders eat a considerable amount of insects in their diets (e.g., 21% for *Callithrix*, see Table 3; and 5-16% reported for *Cebuella*, Ramirez et al. 1977) platyrrhines belonging to this dietary group may still require molars that are capable of breaking and puncturing the hard exoskeleton of insects in order to access the protein inside. This is in contrast with the degenerate, peg-like molars of the honey possum *Tarsipes* (Rosenberg and Richardson 1995; Beck et al. 2022) and the ‘simple’ molars of nectarivorous bats that are decreased in complexity and curvature (measured as OPCR and DNE, López-Aguirre et al. 2022), that, presumably because they are specialised to an almost exclusively liquid diet, lost dental adaptations for other foods.

## Conclusion

We conclude that our freeware pipeline is accurate for widespread use by researchers wishing to carry out their own dental topographic analyses. Our freeware pipeline also provides potential benefits for researchers in institutions or countries that lack funding for proprietary software, but who are nevertheless interested in using DTMs. We also conclude that the pipeline, in combination with the expanded comparative platyrrhine sample of second lower molars and revised dietary classification scheme presented here, is suitable for inferring probable diet of specimens for which direct dietary information is unavailable, such as fossils.

## Statements and Declarations

### Authors’ contributions

Dorien de Vries and Robin Beck conceived the study with assistance from Jean Boubli and Mareike Janiak for the dietary classification scheme. The casts of callitrichid molars were made and μCT scanned by Doug Boyer and Elizabeth St. Clair. Data processing was done by Dorien de Vries with assistance from Emma Ridgway, Elizabeth St. Clair, and Doug Boyer. Analyses were performed by Dorien de Vries. The first draft of the manuscript was written by Dorien de Vries and Robin Beck and all authors commented on later versions of the manuscript. All authors read and approved the final manuscript.

### Competing interests

The authors declare no competing interests.

### Data availability

The datasets generated during and/or analysed during the current study are included in this published article and its supplementary information files. Digital surface meshes of the teeth analysed in this article are available on the Morphosource database (project ID: 000471738) and the original meshes of the 39 callitrichid m2s have the following DOIs: https://doi.org/10.17602/M2/M471790; https://doi.org/10.17602/M2/M471815; https://doi.org/10.17602/M2/M471870; https://doi.org/10.17602/M2/M471878; https://doi.org/10.17602/M2/M471887; https://doi.org/10.17602/M2/M471962; https://doi.org/10.17602/M2/M473213; https://doi.org/10.17602/M2/M473283; https://doi.org/10.17602/M2/M473343; https://doi.org/10.17602/M2/M473386; https://doi.org/10.17602/M2/M473433; https://doi.org/10.17602/M2/M473494; https://doi.org/10.17602/M2/M474623; https://doi.org/10.17602/M2/M474631; https://doi.org/10.17602/M2/M474639; https://doi.org/10.17602/M2/M474647; https://doi.org/10.17602/M2/M474655; https://doi.org/10.17602/M2/M475063; https://doi.org/10.17602/M2/M475103; https://doi.org/10.17602/M2/M475111; https://doi.org/10.17602/M2/M475119; https://doi.org/10.17602/M2/M476460; https://doi.org/10.17602/M2/M476468; https://doi.org/10.17602/M2/M476478; https://doi.org/10.17602/M2/M476492; https://doi.org/10.17602/M2/M476508; https://doi.org/10.17602/M2/M476527; https://doi.org/10.17602/M2/M476552; https://doi.org/10.17602/M2/M476576; https://doi.org/10.17602/M2/M477126; https://doi.org/10.17602/M2/M477134; https://doi.org/10.17602/M2/M477142; https://doi.org/10.17602/M2/M477150; https://doi.org/10.17602/M2/M477158; https://doi.org/10.17602/M2/M477190; https://doi.org/10.17602/M2/M477198; https://doi.org/10.17602/M2/M477206; https://doi.org/10.17602/M2/M477226; https://doi.org/10.17602/M2/M477237.

## Acknowledgements

We thank Tommy Burch for uploading digital surface meshes onto MorphoSource. This research was funded by the Natural Environment Research Council (NERC, NE/T000341/1). Data collection was supported by funds from National Science Foundation (NSF, BCS-1552848 and BCS-1304045) to D.M. Boyer and E. St. Clair.

## Notes

https://www.morphosource.org/projects/000471738?locale=en

### Competing Interest Statement

The authors have declared no competing interest.

