## Supplementary Information for "Comparison of dental topography of marmosets and tamarins (Callitrichidae) to other platyrrhine primates using a novel freeware pipeline"

Supplementary details on dietary classification scheme methods

Grouped under **Fruits**

- Ripe fruit
- Pulp of ripe fruits
- Unripe/green fruit
- 'whole fruits'
- mesocarp (exocarp discarded, seed ingested)
- mesocarp only
- aril
- Exo-and mesocorp/pericarp/perigonium (seeds dropped)
- Exocarp (rest of fruit dropped)
- Juice (chewed fruit dropped)
- Pseudofruit (fleshy pedicel) (fruit dropped)
- Unknown fruit parts
- Figs
- ripe fruits, arils, exudates (single occurrence of this category and in that case was only 1%)
- fruits including fruits with small seeds
- Inflorescence
- Flowers, nectar, pollen
- Flower buds
- Nectar from flowers
- Flowers (most time they only drank the nectar
- blossoms
- ‘reproductive parts of *Brosimum alicastrum* (include female flowers, very young fruit, and perhaps male flowers, in unknown proportions)’
- ‘Nectar from flowers’
- ‘Flowers, most time they only drank the nectar’
- Fungi/Fungus

After careful consideration of the data, we grouped ‘Fruit’, ‘Flowers’, and ‘Fungi’ together into one ‘Fruit’ category. ‘Flowers’ were grouped with ‘Fruit’ rather than with

‘Leaves’ based on the functional properties of flowers, which are believed to be softer than those of leaves and structurally more similar to fruits than to leaves. The ‘Fungi’ category was added to ‘Fruits’ as leaving ‘Fungi’ as its own category resulted in *Callimico* to be grouped on its own, separate from all other platyrrhine genera in the UPGMA tree. As *Callimico* would have been the only representative of the ‘Fungivore’ category, we chose not to have this dietary category on its own as we would not be able to test whether the fungivore dental shape was purely phylogenetic, or whether it was driven by the fungivore diet. We thus created dietary categories that were represented by at least two genera.

Grouped under **Leaves**

- Young leaves
- Mature leaves
- Petioles
- Petioles and pulvinus
- Shoots
- Green stems
- Leaf buds
- Orchid pseudobulb
- Leaves, buds, shoots and other vegetational parts of plants
- Young leaves, shoots, petioles
- Aerial roots / tips of aerial roots
- ‘leaves, petioles, bromeliads’
- Bromeliad leaves
- Meristems
- vegetative material (pith, young leaves, etc.)
- tendrils
- leaves, stems, roots, and flowers
- Legumes
- Pods
- ‘petioles & twigs’
- ‘twigs, stems, bark’
- Twigs
- ‘Bark, bamboo, buds, and ferns’
- ‘leaves, petiole bases, tree bark, and unidentifiable plant parts’
- Bark
- Wood (alive or decaying)
- Moss
- Decaying wood/rotten sheaths of *Attalea regia*
- Roots
- Piths
- Loranthaceae (showy mistletoe, family of flowering plants). Referred to as ‘parasite’ grouped under ‘fruits/seeds’ by Norconk 1996

Grouped under **Animal matter**

- Arthropods
- Insects
- Insect eggs
- Caterpillars
- Termites
- Insects and spiders
- caterpillars, larvae, etc.
- insects, insect eggs, larvae, lizards
- frogs, insects, ants
- vertebrates

Grouped under **Seeds**

- Seeds (Includes fruit, e.g., whole fruit, from which seeds were specified as having been eaten)
- Palm nuts
- Other seeds and nuts
- Seeds of nuts and drupes (*Sapotaceae, Chrysobalanaceae, Connaraceae, Moraceae, Sapindaceae, Anacardiaceae. Rubiaceae*).
- Winged seeds (*Bignoniaceae, Polygalaceae. Combretaceae, Tiliaceae*)
- Legume seeds (*Caesalpiniaceae, Papilionaceae. Mimosaceae*)

Grouped under **Other**

- Unidentified / unspecified
- arboreal termite nest
- Termitaria or Termitaria earth
- ‘invertebrates, leaf galls, bark, and fungi’
- ‘Bark and undetermined items’
- ‘wood, soil, termitaria’ – exclude (which means up the ‘other’ category

**Excluded and increased each of the other categories by x = % excluded category/remaining categories**

- corn from surrounding plantations – Exclude
- ‘soil from termitaria nest, fungi, and a frog’ – Exclude
- fluids of unripe palm nuts (Astrocaryum) - Exclude
- Buds (also referred to as “broto” in Spanish paper). If no further detail was provided this was excluded

**Table S1** Raw dietary data per genus showing the breakdown of the combined 'Fruits, Flowers, and Fungi' category. These raw data are ordered per dietary cluster following the UPGMA outcome (see main text Figure 2 and Table 3)

| Genus | Fruits | | | | Leaves | Seeds | Animal matter | Exudates | Other |
|---|---|---|---|---|---|---|---|---|---|
| | Total | Fruits | Flowers | Fungi | | | | | |
| **Exudate feeder** | | | | | | | | | |
| *Callithrix* | 0.28 | 0.261 | 0.003 | 0.016 | 0 | 0 | 0.21 | 0.52 | <0.01 |
| **Seed feeder** | | | | | | | | | |
| *Cheracebus* | 0.467 | 0.447 | 0.02 | 0 | 0.09 | 0.32 | 0.09 | 0 | 0 |
| *Chiropotes* | 0.447 | 0.418 | 0.029 | 0 | 0.04 | 0.45 | 0.02 | 0 | 0.04 |
| *Cacajao* | 0.192 | 0.137 | 0.056 | 0 | 0.037 | 0.69 | 0.04 | 0 | 0 |
| **Folivore** | | | | | | | | | |
| *Alouatta* | 0.404 | 0.316 | 0.088 | 0 | 0.57 | <0.01 | <0.01 | 0 | 0.01 |
| *Brachyteles* | 0.464 | 0.373 | 0.091 | 0 | 0.52 | 0 | 0 | 0 | 0 |
| **Frugivore** | | | | | | | | | |
| *Ateles* | 0.833 | 0.799 | 0.034 | <0.01 | 0.12 | 0.02 | <0.01 | 0 | 0.02 |
| *Callicebus* | 0.704 | 0.699 | 0.005 | 0 | 0.16 | 0.13 | 0 | 0 | <0.01 |
| *Aotus* | 0.638 | 0.605 | 0.033 | 0 | 0.24 | 0 | 0.03 | 0 | 0 |
| *Lagothrix* | 0.693 | 0.677 | 0.016 | <0.01 | 0.12 | 0.05 | 0.12 | 0 | 0.01 |
| *Plecturocebus* | 0.581 | 0.579 | 0.002 | 0 | 0.28 | 0 | 0.10 | 0 | <0.01 |

| | | | | | | | | | |
|---|---|---|---|---|---|---|---|---|---|
| *Pithecia* | 0.602 | 0.557 | 0.045 | 0 | 0.17 | 0.21 | 0.02 | 0.01 | <0.01 |
| **Frugivore-Insectivore** | | | | | | | | | |
| *Callimico* | 0.58 | 0.29 | 0 | 0.29 | 0 | 0 | 0.34 | 0.01 | 0.07 |
| *Cebus* | 0.58 | 0.565 | 0.015 | 0 | 0.06 | 0.03 | 0.27 | 0 | 0.03 |
| *Leontocebus* | 0.56 | 0.53 | 0.03 | 0 | 0 | 0 | 0.27 | 0.12 | 0.01 |
| *Leontopithecus* | 0.79 | 0.68 | 0.11 | 0 | 0 | 0 | 0.13 | 0.08 | <0.01 |
| *Saguinus* | 0.649 | 0.612 | 0.038 | 0 | 0 | 0 | 0.24 | 0.07 | 0.02 |
| *Saimiri* | 0.637 | 0.602 | 0.035 | 0 | 0 | 0.01 | 0.31 | 0 | 0 |
| *Sapajus* | 0.439 | 0.408 | 0.031 | 0 | 0.25 | 0.05 | 0.27 | 0 | <0.01 |

**Table S2** Total platyrrhine sample: 150 specimens, of which 111 were previously studied by Winchester et al. (2014), and 39 are newly added callitrichid specimens (highlighted in bold)

| Taxon | Specimen number |
|---|---|
| *Alouatta palliata* | usnm171063 |
| *Alouatta palliata* | usnm284782 |
| *Alouatta palliata* | usnm290601 |
| *Alouatta seniculus* | SBU_NAl13 |
| *Alouatta seniculus* | usnm123517 |
| *Alouatta seniculus* | usnm281658 |
| *Alouatta seniculus* | usnm281673 |
| *Alouatta seniculus* | usnm281741 |
| *Alouatta seniculus* | usnm281751 |
| *Alouatta seniculus* | usnm281758 |
| *Aotus azarae* | AMNH211460 |
| *Aotus azarae* | AMNH211465 |
| *Aotus nigriceps* | AMNH147472 |
| *Aotus nigriceps* | AMNH67246 |
| *Aotus nigriceps* | AMNH75996 |
| *Aotus nigriceps* | AMNH75999 |
| *Aotus nigriceps* | AMNH76002 |
| *Aotus nigriceps* | AMNH92804 |
| *Aotus nigriceps* | AMNH92809 |
| *Aotus nigriceps* | usnm364486 |
| *Ateles belzebuth* | AMNH71787 |
| *Ateles belzebuth* | AMNH76882 |
| *Ateles belzebuth* | AMNH67102 |
| *Ateles belzebuth* | usnm406674 |
| *Ateles belzebuth* | usnm406675 |
| *Ateles geoffroyi* | MCZ34320 |
| *Ateles geoffroyi* | MCZ5344 |
| *Ateles geoffroyi* | usnm336204 |

| | |
|---|---|
| *Ateles paniscus* | MCZ31759 |
| *Brachyteles arachnoides* | MCZ5070 |
| *Brachyteles arachnoides* | MN-Rio106 |
| *Brachyteles arachnoides* | MN-Rio24104 |
| *Brachyteles arachnoides* | MN-Rio24114 |
| *Brachyteles arachnoides* | MN-Rio2718 |
| *Brachyteles arachnoides* | MN-Rio30191 |
| *Brachyteles arachnoides* | MN-Rio526 |
| *Brachyteles arachnoides* | MN-Rio6699 |
| *Brachyteles arachnoides* | MN-Rio7724 |
| *Brachyteles arachnoides* | MN-Rio8513 |
| *Cacajao calvus* | AMNH73720 |
| *Cacajao calvus* | AMNH76391 |
| *Cacajao calvus* | AMNH76648 |
| *Cacajao calvus* | AMNH98397 |
| *Cacajao calvus_rubicundus* | SBU_NCj01 |
| *Cacajao melanocephalus* | AMNH78566 |
| *Cacajao melanocephalus* | AMNH78569 |
| *Cacajao melanocephalus* | usnm256215 |
| *Cacajao melanocephalus* | usnm406423 |
| *Plecturocebus donacophilus* | AMNH211491 |
| *Plecturocebus donacophilus* | AMNH211494 |
| *Plecturocebus donacophilus* | AMNH40834 |
| *Plecturocebus moloch* | AMNH94972 |
| *Plecturocebus moloch* | AMNH94979 |
| *Cheracebus torquatus* | AMNH76861 |
| *Cheracebus torquatus* | AMNH77300 |
| *Cheracebus torquatus* | AMNH78466 |
| *Cheracebus torquatus* | AMNH78473 |
| *Cheracebus torquatus* | AMNH78580 |
| ***Callimico goeldii*** | **USNM303322** |

| | |
|---|---|
| ***Callimico goeldii*** | **USNM303323** |
| ***Callimico goeldii*** | **USNM395455** |
| ***Callimico goeldii*** | **USNM399073** |
| ***Callimico goeldii*** | **USNM464993** |
| ***Callimico goeldii*** | **USNM575153** |
| ***Callimico goeldii*** | **USNM582737** |
| ***Callithrix jacchus penicillata*** | **MCZ37821** |
| ***Callithrix jacchus penicillata*** | **USNM259427** |
| ***Cebuella pygmaea*** | **USNM336305** |
| ***Cebuella pygmaea*** | **USNM336307** |
| ***Cebuella pygmaea*** | **USNM336318** |
| ***Cebuella pygmaea*** | **USNM336320** |
| *Sapajus apella* | AMNH239863 |
| *Sapajus apella* | AMNH239864 |
| *Sapajus apella* | AMNH78501 |
| *Sapajus apella* | usnm296634 |
| *Sapajus apella* | usnm361019 |
| *Sapajus apella* | usnm461384 |
| *Cebus capucinus* | usnm291123 |
| *Cebus capucinus* | usnm291128 |
| *Cebus capucinus* | usnm464845 |
| *Chiropotes albinasus* | AMNH461707 |
| *Chiropotes albinasus* | AMNH545874 |
| *Chiropotes albinasus* | AMNH95302 |
| *Chiropotes albinasus* | MCZ31701 |
| *Chiropotes albinasus* | usnm543357 |
| *Chiropotes satanas* | usnm338962 |
| *Chiropotes satanas* | usnm406583 |
| *Chiropotes satanas* | usnm406593 |
| *Chiropotes satanas* | usnm546263 |
| *Chiropotes satanas* | usnm546264 |

| | |
|---|---|
| *Chiropotes satanas* | usnm549519 |
| *Lagothrix lagotricha poepiggi* | AMNH188142 |
| *Lagothrix lagotricha poepiggi* | AMNH71776 |
| *Lagothrix lagotricha poepiggi* | AMNH71780 |
| *Lagothrix lagotricha poepiggi* | AMNH76393 |
| *Lagothrix lagotricha poepiggi* | AMNH98332 |
| *Lagothrix lagotricha* | usnm545879 |
| *Lagothrix lagotricha* | usnm545887 |
| *Lagothrix lagotricha* | usnm545890 |
| ***Leontopithecus rosalia*** | **MCZ3939** |
| ***Leontopithecus rosalia*** | **USNM337333** |
| ***Leontopithecus rosalia*** | **USNM337334** |
| ***Leontopithecus rosalia*** | **USNM541408** |
| ***Leontopithecus rosalia*** | **USNM545068** |
| ***Leontopithecus rosalia*** | **USNM582752** |
| ***Mico argentata*** | **USNM239457** |
| ***Mico argentata*** | **USNM239458** |
| ***Mico argentata*** | **USNM239459** |
| ***Mico argentata*** | **USNM239461** |
| ***Mico argentata*** | **USNM461725** |
| ***Mico argentata*** | **USNM555657** |
| *Pithecia monachus* | AMNH6412 |
| *Pithecia monachus* | MCS30720 |
| *Pithecia monachus* | usnm461919 |
| *Pithecia monachus* | usnm545891 |
| *Pithecia pithecia* | usnm374744 |
| *Pithecia pithecia* | usnm374745 |
| *Pithecia pithecia* | usnm374746 |
| *Pithecia pithecia* | usnm374756 |
| *Pithecia pithecia* | usnm374759 |
| *Pithecia pithecia* | usnm374767 |

| | |
|---|---|
| ***Leontocebus fuscicollis*** | **AMNH73389** |
| ***Leontocebus fuscicollis*** | **AMNH73392** |
| ***Leontocebus fuscicollis*** | **AMNH73393** |
| ***Leontocebus fuscicollis*** | **AMNH73736** |
| ***Saguinus geoffroyi*** | **USNM335455** |
| ***Saguinus midas*** | **USNM393800** |
| ***Saguinus midas*** | **USNM393802** |
| ***Saguinus midas*** | **USNM393810** |
| ***Saguinus midas*** | **USNM549521** |
| ***Saguinus mystax*** | **USNM397870** |
| ***Saguinus mystax*** | **USNM397877** |
| ***Saguinus oedipus*** | **USNM336297** |
| ***Saguinus oedipus*** | **USNM501096** |
| ***Saguinus oedipus*** | **USNM501106** |
| *Saimiri boliviensis* | AMNH38792 |
| *Saimiri boliviensis* | AMNH76003 |
| *Saimiri boliviensis* | AMNH76583 |
| *Saimiri boliviensis* | AMNH76586 |
| *Saimiri boliviensis* | AMNH98272 |
| *Saimiri boliviensis* | usnm364497 |
| *Saimiri boliviensis* | usnm396265 |
| *Saimiri sciureus* | usnm518547 |
| *Saimiri sciureus* | usnm545893 |
| *Saimiri sciureus* | usnm546267 |
| The following five specimens were only included in the validation test of the new pipeline when the original sample of Winchester et al. (2014) was replicated. | |
| *Ateles belzebuth* | usnm241384 |
| *Cacajao calvus* | AMNH98316 |
| *Cebus capucinus* | usnm291133 |
| *Lagothrix lagotricha poepiggi* | AMNH71767 |

| | |
|---|---|
| *Lagothrix lagotricha* | usnm545878 |

**Table S3** One-way ANOVA results. Games-Howell post hoc tests were applied when the one-way ANOVA was significant (see Table S4)

| **Variable** | **df** | ***F*** | ***P*** |
|---|---|---|---|
| ln(2D area) | 4 | 101.5 | <0.01 |
| OPCR | 4 | 87.5 | <0.01 |
| RFI | 4 | 16.11 | <0.01 |
| Slope | 4 | 11.32 | <0.01 |
| DNE | 4 | 0.127 | 0.97 |
| Convex DNE | 4 | 1.128 | 0.35 |

**Table S4** P-values of pairwise comparisons of Games-Howell post hoc tests applied to the significantly different dental topographic variables (as identified through one-way ANOVAs, see Table S3). Significant (<0.05) pairwise comparisons are in bold. Fo = Folivory, Fr-In = Frugivory-Insectivory, Fr = Frugivory, Se = Seed eating, Ex = Exudate feeding. See SM for a list of degrees of freedom and the F values

| Variables: | ln(2D area) | OPCR | RFI | Slope |
|---|---|---|---|---|
| Dietary categories compared: | *P* | *P* | *P* | *P* |
| Ex : Fo | **<0.01** | **<0.01** | **0.043** | >0.99 |
| Ex : Fr-In | **<0.01** | 0.096 | >0.99 | 0.99 |
| Ex : Fr | **<0.01** | 0.11 | 0.46 | 0.75 |
| Ex : Se | **<0.01** | **<0.01** | **0.005** | **<0.01** |
| Fo : Fr-In | **<0.01** | **<0.01** | **0.013** | 0.991 |
| Fo : Fr | **<0.01** | **<0.01** | **<0.01** | 0.34 |
| Fo : Se | **<0.01** | **<0.01** | **<0.01** | **<0.01** |
| Fr-In : Fr | **<0.01** | **<0.01** | 0.38 | 0.33 |
| Fr-In : Se | **<0.01** | **<0.01** | **<0.01** | **<0.01** |
| Fr : Se | 0.77 | **<0.01** | **<0.01** | **<0.01** |

**Table S5** Average, ranges, and variance of dental metrics per dietary group of the preferred dietary scheme (edited UPGMA scheme) and total sample (n = 145)

| Metric | Dietary group (n) | mean | range | variance |
|---|---|---|---|---|
| **ln(2D area)** | Folivory (20) | 3.69 | 3.44-3.88 | 0.02 |
| | Frugivory (37) | 2.7 | 2.11-3.48 | 0.25 |
| | Frugivory-Insectivory (46) | 1.82 | 0.97-3.07 | 0.37 |
| | Seed eating (30) | 2.60 | 2.26-2.93 | 0.03 |
| | Exudate feeding (12) | 0.92 | 0.51-1.33 | 0.07 |
| **OPCR** | Folivory (20) | 112.22 | 88.62-147.25 | 292.34 |
| | Frugivory (37) | 169.8 | 111.25-235.62 | 845.43 |
| | Frugivory-Insectivory (46) | 138.7 | 102.38-199 | 577.33 |
| | Seed eating (30) | 234.8 | 189.75-294.50 | 1029.78 |
| | Exudate feeding (12) | 153.5 | 126.38-176.25 | 234.45 |
| **RFI** | Folivory (20) | 0.53 | 0.46-0.58 | <0.01 |
| | Frugivory (37) | 0.48 | 0.43-0.54 | <0.01 |
| | Frugivory-Insectivory (46) | 0.50 | 0.41-0.57 | <0.01 |
| | Seed eating (30) | 0.46 | 0.43-0.51 | <0.01 |
| | Exudate feeding (12) | 0.50 | 0.43-0.53 | <0.01 |
| **Slope** | Folivory (20) | 70.90 | 68.65-73.57 | 2.22 |
| | Frugivory (37) | 70.09 | 65.77-72.69 | 2.58 |
| | Frugivory-Insectivory (46) | 71.12 | 63.77-76.69 | 10.29 |
| | Seed eating (30) | 67.82 | 64.90-71.25 | 1.95 |
| | Exudate feeding (12) | 70.78 | 66.63-73.28 | 3.04 |
| **DNE** | Folivory (20) | 240.3 | 177.93-299.74 | 997.9 |

| | | | | |
|---|---|---|---|---|
| | Frugivory (37) | 246.1 | 192.84-398.37 | 2244.51 |
| | Frugivory-Insectivory (46) | 242.8 | 137.99-306.75 | 1866.5 |
| | Seed eating (30) | 240.1 | 204.07-305.84 | 710.51 |
| | Exudate feeding (12) | 242.1 | 208.7-271.5 | 330.19 |
| **Convex DNE** | Folivory (20) | 176.0 | 130.11-218.24 | 650.29 |
| | Frugivory (37) | 181.4 | 143.18-268.84 | 734.04 |
| | Frugivory-Insectivory (46) | 186.4 | 115.76-238.78 | 1001.91 |
| | Seed eating (30) | 175.8 | 155.95-205.99 | 183.3 |
| | Exudate feeding (12) | 186.2 | 159.3-213.26 | 206.1 |

**Table S6** Results shown for discriminant function analysis using the non-callitrichid platyrrhine sample (n = 145) and topography + size variables (95.28% overall success). Classification accuracy shown as a breakdown of the correctly classified percentage and misclassified percentage per dietary category

| | Classified as: | | | |
|---|---|---|---|---|
| Assigned diet | Folivory | Frugivory-Insectivory | Frugivory | Seed eating |
| Folivory | 0.95 | 0 | 0.05 | 0 |
| Frugivory-Insectivory | 0 | 0.89 | 0.05 | 0.05 |
| Frugivory | 0 | 0 | 1 | 0 |
| Seed eating | 0 | 0 | 0.07 | 0.93 |

**Table S7** Results shown for discriminant function analysis using the entire platyrrhine sample (n = 145) and topography + size variables (90.34% overall success). Classification accuracy shown as a breakdown of the correctly classified percentage and misclassified percentage per dietary category

| | Classified as: | | | | |
|---|---|---|---|---|---|
| Assigned diet: | Folivory | Frugivory-Insectivory | Frugivory | Seed eating | Exudate feeding |
| Folivory | 0.95 | 0 | 0.05 | 0 | 0 |
| Frugivory-Insectivory | 0 | 0.85 | 0.07 | 0.02 | 0.07 |
| Frugivory | 0 | 0.03 | 0.97 | 0 | 0 |
| Seed eating | 0 | 0 | 0.07 | 0.93 | 0 |
| Exudate feeding | 0 | 0.25 | 0 | 0 | 0.75 |
